# Spatial Orchestration of Skin Fibrosis by a CD8+ T cell-Myofibroblast Axis

**DOI:** 10.64898/2026.06.17.732787

**Authors:** Ian C. Boothby, Thomas C. Gan, Vrinda Johri, Emily Flynn, Maha Kazmi, Lenka Maliskova, Saba Shaikh, Shivaram Yellamilli, Gabriela K. Fragiadakis, Isaac Neuhaus, Walter Eckalbar, Jarish N. Cohen, Alexis J. Combes, Anna Haemel, Michael D. Rosenblum, Maxime J. Kinet

## Abstract

Fibrosing skin diseases are highly morbid conditions with diverse clinical and histopathologic features. Prior work, primarily in systemic sclerosis (SSc), has yielded mixed data regarding the immune drivers of fibrosis, as well as the identity and spatial localization of pro-fibrotic fibroblast cell subsets. Here, we focus on morphea and eosinophilic fasciitis (EF), which cause more acutely inflammatory skin fibrosis. Using multimodal single-nucleus and spatial transcriptomics, we find that effector CD8+ T cells are highly enriched in fibrotic skin. These cells are particularly abundant in inflammatory tissue domains bordering fibrotic stroma, which are marked by expression of interferon-γ stimulated genes. Inflammatory domains feature a loss of local homeostatic fibroblast populations and replacement with ADAM12-expressing inflammatory fibroblasts and myofibroblasts, which co-localize closely with CD8+ T cells. All subtypes of morphea featured similar patterns of CD8+ T-cell-associated fibro-inflammatory zonation and fibroblast transformation, suggesting that shared mechanisms can drive fibrosis across stromal compartments of skin. We apply these findings to a large publicly available scleroderma dataset and find that similar processes occur in SSc. Mechanistically, ablation of CD8+ T cells in mice ameliorates bleomycin-driven inflammation and fibrosis, as does fibroblast-intrinsic abrogation of IFN-γ signaling. These data establish CD8+ T cell-driven fibrogenesis as a key feature of fibrosing skin diseases and raise the prospect of targeting CD8+ T cells in autoimmune fibrosis more broadly.

**One sentence summary:** Spatial profiling of morphea-spectrum diseases reveals CD8 T cells as key drivers of fibrosis through fibroblast IFN-γ signaling.

## INTRODUCTION

Fibrosis, the pathological replacement of normal tissue architecture with extracellular matrix secreted by activated fibroblasts, underlies a diverse array of diseases affecting many organ systems. Morphea-spectrum disorders (MSD), including eosinophilic fasciitis (EF), are a spectrum of fibroinflammatory skin diseases that cause significant morbidity through loss of normal skin mobility, elasticity, and wound healing (*1*). Several subtypes of MSD are defined by the depth and distribution of inflammation and fibrosis. Superficial morphea affects the papillary dermis; plaque and generalized morphea extend into the reticular dermis and subcutaneous adipose tissue; and deep morphea involves the deep dermis, subcutaneous fat, and fascia (*1*). EF, characterized by eosinophil-rich inflammation and fibrosis of the subcutaneous fat and deep fascia, occupies the most severe end of this clinicopathologic spectrum, often involving entire limbs (*2*). The histopathology of morphea can closely resemble systemic sclerosis (SSc), a distinct systemic fibroinflammatory autoimmune disease, hinting at shared mechanisms across cutaneous fibrosis. It is unknown if these heterogeneous presentations of MSDs share the same inflammatory and fibroblastic drivers. Furthermore, mechanisms that initiate fibroinflammatory reactions in a single organ’s different histologic compartments remain incompletely understood, and we lack effective disease-modifying therapies (*3*).

In MSD and SSc, innate and adaptive immune activation precede and accompany fibroblast activation (*4*, *5*). CD4+ T helper subsets—particularly T_h_2 and T_h_17 cells—have been proposed to drive profibrotic cytokine cascades, whereas type 1 immunity has historically been thought to limit fibrotic responses (*6*). Indeed, interferon-gamma (IFN-γ), the defining cytokine of T_h_1 and CD8+ cytotoxic T cell responses, was long regarded as an antifibrotic mediator. Early studies demonstrated that IFN-γ dampens fibroblast TGF-β signaling and collagen synthesis, as well as tissue fibrosis, in several organs (*7–19*). However, clinical trials of IFN-γ in SSc and MSD showed mixed results (*20–31*). Together, these observations raise the possibility that the antifibrotic effects of IFN-γ described in earlier reductionist systems may not generalize across disease contexts and that exogenous and endogenously produced IFN-γ may have different effects. Moreover, the role of cytotoxic CD8+ T cells, a dominant cellular source of IFN-γ, has received comparatively little direct mechanistic attention in fibrotic disease, and relevant cellular targets of IFN-γ in skin fibrosis have not been conclusively established.

Here, we apply single-nuclear RNA-sequencing and spatial transcriptomics to comprehensively define the immune and stromal landscapes across MSD. We identify the replacement of homeostatic fibroblasts by inflammatory and pro-fibrotic fibroblasts within disease-subtype-specific, depth-stratified zones of fibroinflammation. We demonstrate that inflammatory fibroblasts in these zones exhibit a prominent IFN-γ-driven transcriptional response. Unexpectedly, we find that CD8+ T cells are the most expanded immune population across disease subtypes and that these cells colocalize with fibroinflammatory tracts. Importantly, we extend these findings to an existing SSc dataset. Lastly, we evaluate the functional importance of CD8-fibroblast crosstalk in a murine model of skin fibrosis, demonstrating that depleting CD8+ T cells attenuates fibrosis, as does abrogating IFN-γ signaling within fibroblasts. Collectively, these data establish a CD8+ T cell–IFN-γ–fibroblast axis as a driver of cutaneous fibrosis and nominate this pathway as a tractable therapeutic target across MSD and scleroderma.

## RESULTS

### Fibrosing skin is zonated into inflammatory and fibrotic tissue domains

In MSD, new inflammatory lesions often appear acutely and progress to fibrosis over the course of weeks to months, in contrast to the more chronic progression of fibrosis in diseases like SSc. Areas of inflammation and fibrosis are often clinically visible in the same lesion, enabling direct observation of fibroinflammatory progression within a single biopsy. We collected a cohort of 21 paraffin-embedded skin biopsies from patients with active morphea-spectrum lesions and 11 healthy controls **(Table S1)**. We performed single-nuclear RNA sequencing (snRNAseq) on all samples as well as Visium and Xenium spatial transcriptomics (ST) on a subset of these same samples (**Fig. 1A, S1A-J).** We additionally performed label transfer to generate unified annotations across snRNAseq and Xenium data, enabling direct comparison **(Fig. S1K)**.

**Fig. 1.**
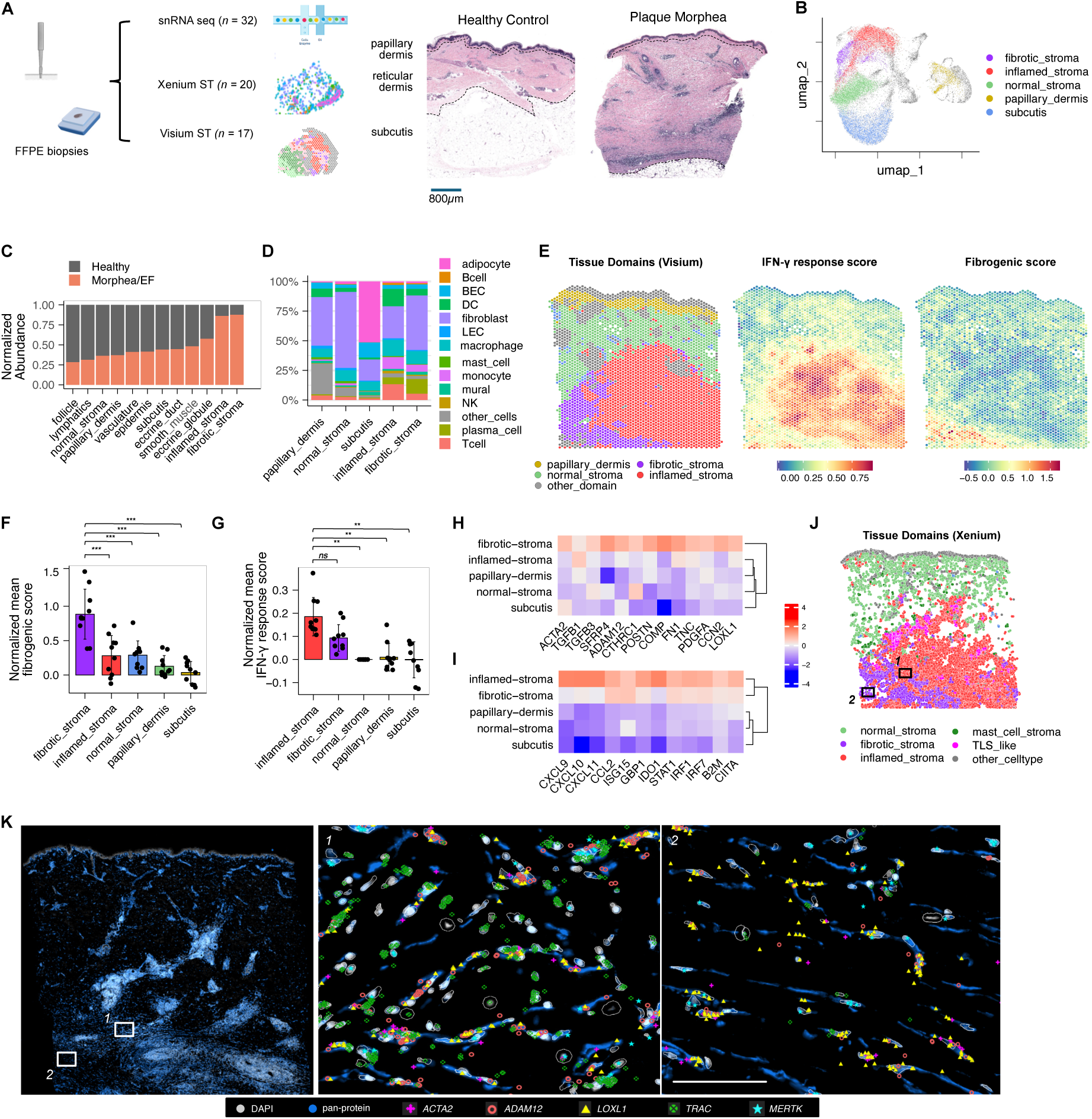
Fibrosing skin is zonated into inflammatory and fibrotic domains. **(A)** Schematic of experimental design. FFPE-formalin-fixed paraffin-embeded tissue; snRNAseq - single nuclear RNA sequencing; ST-spatial transcriptomics. Representative micrographs of sections from healthy skin and plaque morphea stained with hematoxylin and eosin (H&E). Dotted lines outline approximate boundaries between relevant skin layers. **(B)** Uniform Manifold Approximation and Projection (UMAP) plot of stroma-predominant tissue domains in Visium data. **(C)** Comparative normalized abundance of each tissue domain in healthy and fibrosis samples. **(D)** Frequencies of predicted immune and stromal cell types in each Visium tissue domain using snRNAseq data as a reference atlas. **(E)** Representative tissue domain distribution and localization of IFN-γ response scores/fibrogenic scores in an eosinophilic fasciitis (EF) sample. **(F-G)** Expression of fibrosis-associated (F) and IFN-γ stimulated genes (G) in each tissue domain. **(H-I)** Per-sample fibrogenic (H) and IFN-γ response (I) scores in Visium data, normalized to the normal stroma tissue domain. Statistics calculated using one-way ANOVA, *p* = 1.85 × 10^−10^ (G) and 1.68 × 10^−3^ (H)**. (J)** Tissue domains identified in Xenium data of an EF sample. **(K)** Representative image of EF tissue with insets demonstrating inflammatory stroma (*1*) and fibrotic stroma (*2*). Inset scale bar: 100µm. *All plots:* \**p* < 0.05; \*\**p* < 0.05; \*\*\**p* < 0.005; *ns*, not significant.

Using Leiden-based clustering of Visium data, we observed spatial demarcation of lesional skin into inflammatory and fibrotic domains, which were significantly enriched in morphea and EF samples (**Fig. 1B-C, Fig. S2A).** Unaffected dermis and subcutaneous tissue was primarily composed of fibroblasts, adipocytes, and other stromal cells (**Fig**. **1D**), whereas inflammatory domains, and, to a lesser extent, fibrotic domains, were infiltrated by T cells, monocytes, and dendritic cells **(Fig. 1D).** At the molecular level, fibrotic domains were enriched for extracellular matrix-related transcriptional programs and strongly expressed activation- and fibrosis-related genes such as *POSTN*, *COMP,* and *ADAM12* (**Fig. 1E, F, H; Fig. S2B-C)**. Both inflamed and fibrotic stroma were enriched in interferon gamma (IFN-γ) stimulated signatures, with particular enrichment in inflamed stroma **(Fig. 1E, G, I; Fig. S2B-C).** IFN-α-stimulated gene signatures were also upregulated in both inflamed and fibrotic domains, although to a lesser degree than for IFN-γ **(Fig. S2B).**

We achieved similar results using an orthogonal approach to identify tissue domains by constructing Tessera tiles from neighboring single cells in Xenium data **(Fig. S2D,E)** (*32*). Higher spatial resolution enabled identification of finer tissue domains such as tertiary lymphoid-like (TLS-like) structures, although the papillary dermis was not well-distinguished from the epidermis **(Fig. S2D)**. Again, zonation of inflamed and fibrotic stromal domains was seen **(Fig. 1J)**, with distinct inflammatory infiltrates **(Fig. S2E)** and patterns of inflammatory and pro-fibrotic gene expression, respectively **(Fig S2E-F).** At single-cell spatial resolution, we were able to observe dense infiltrates of T cells, macrophages, and fibroblasts in inflamed stroma, whereas fibrotic stroma was comparatively more hypocellular and dominated by *ACTA2^+^* myofibroblasts **(Fig. 1K).**

Next, we compared the spatial distribution of inflammation and fibrosis across different MSD subtypes, which feature fibrosis in different microanatomic compartments of skin (papillary dermis, reticular dermis, or subcutis) **(Fig. 2A)**. In superficial morphea (SM), which affects the uppermost layers of skin, the papillary and upper reticular dermis were primarily affected. Plaque and generalized morphea (PM/GM) specimens featured fibrosis of the entire thickness of the reticular dermis with occasional involvement of the upper subcutis. In morphea profunda and EF (MP/EF), linearly-arranged strata of inflamed and fibrotic tissue domains could be seen in the subcutis due to inflammation of the fibrous septae that separate lobules of adipocytes in skin **(Fig. 1J,K**; **Fig. 2A).** The depth from the epidermal surface of these pathologic domains stratified according to MSD subtype **(Fig 2B, C)**. Extending classical histopathologic descriptions of disease, these data demonstrate that a similar fibro-inflammatory reaction, at the level of coarse gene expression, occurs across MSD subtypes spanning the full depth of the skin and subcutis.

**Fig. 2.**
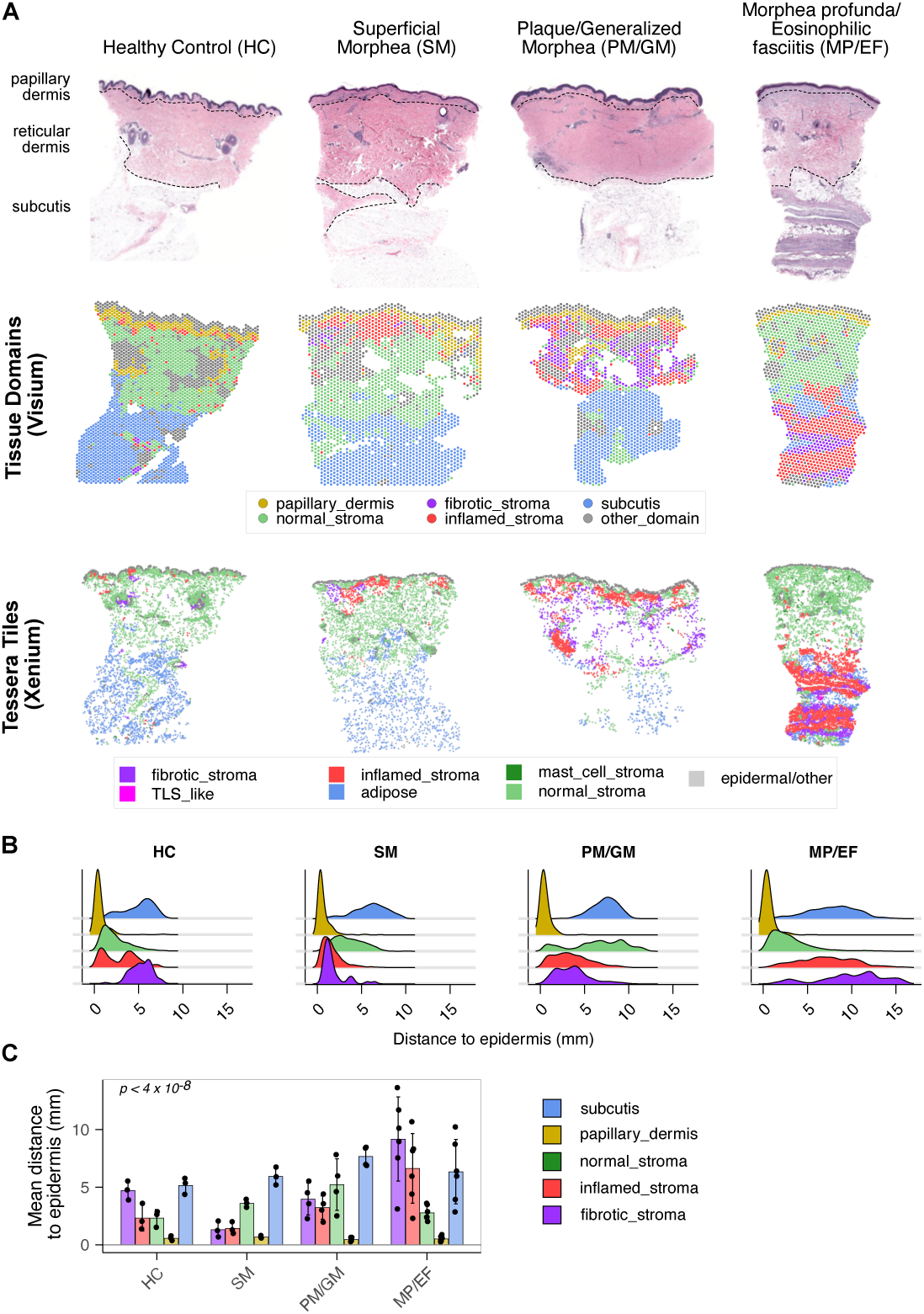
Fibro-inflammatory patterning is conserved across disease subtypes and cutaneous tissue compartments. **(A)** Representative H&E images, tissue domain distribution, and fibroblast subset localization in subtypes of morphea and eosinophilic fasciitis, grouped by depth of involvement. **(B-C)** Depth of involvement of fibrosis and inflammation across morphea and EF subtypes. Statistics in (C) calculated using a linear mixed-effects model with disease subtype and tissue domain as main effects; Chi-square p-value for the interaction term is reported.

### A full-thickness spatial atlas of skin fibroblasts

We next focused on the fibroblast subtypes present in healthy, inflamed, and fibrotic stromal domains. As structural cells, fibroblasts’ identity is determined in part by their anatomic compartment within tissues. We therefore used careful spatial analysis to annotate transcriptional clusters of cell types in the snRNAseq data **(Fig. 3A, Fig. S3A)**. Our results agree with a recent atlas of skin fibroblasts (*33*). In the papillary dermis, papillary fibroblasts express the Wnt pathway related genes *RSPO1*, *AXIN2*, and *APCDD1* **(Fig. 3B-D; Fig. S3B-F)**. The reticular dermis contains numerous reticular fibroblasts marked by *PI16, LGR5,* and *CD34* **(Fig. 3B, C)**. These fibroblasts could also be seen extending into the interlobular fibrous septae of the subcutis **(Fig. S3B, C).**

**Fig. 3.**
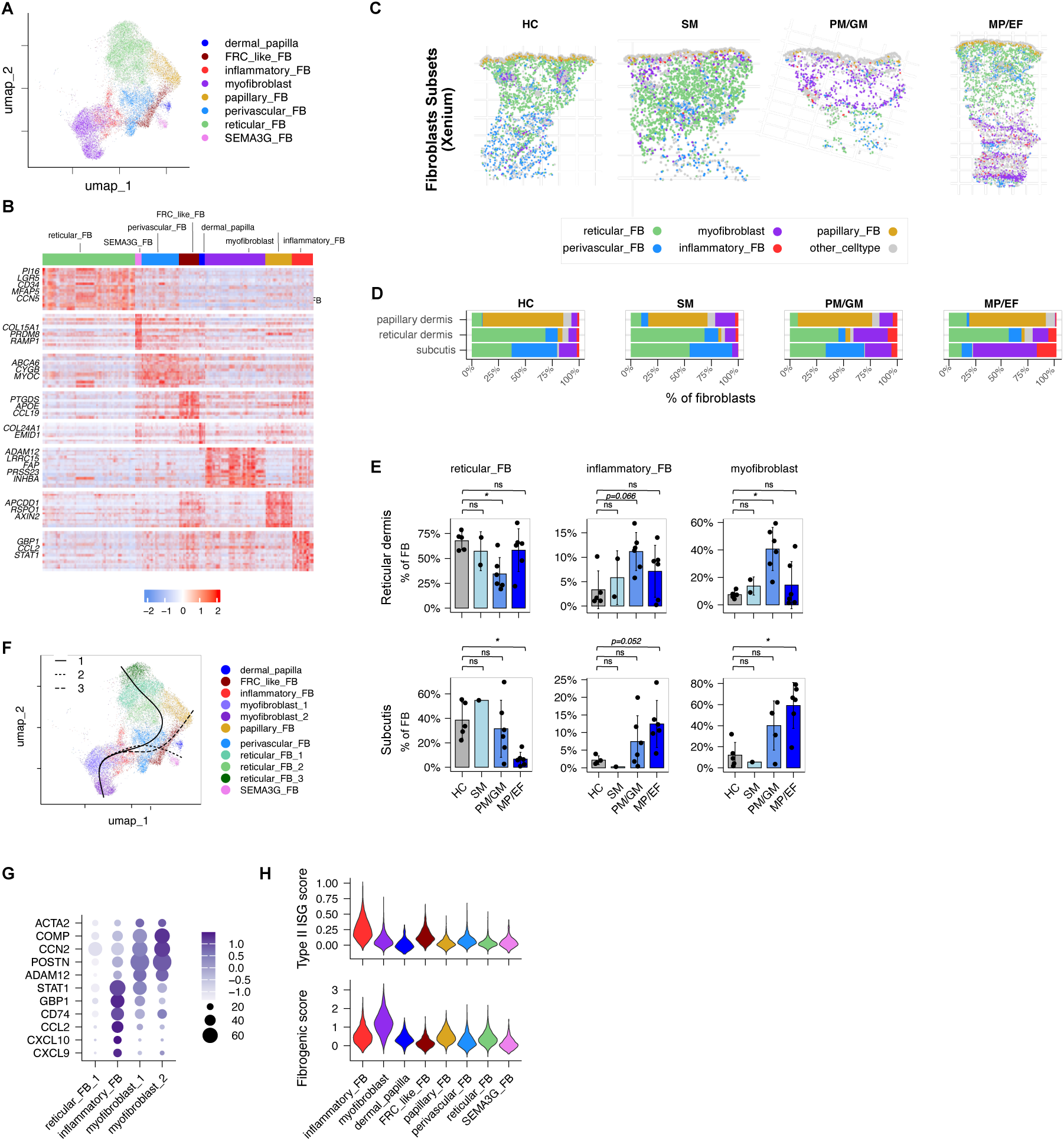
Local homeostatic fibroblast subsets are replaced by interferon-stimulated inflammatory fibroblasts and *ADAM12+* myofibroblasts in fibrosing skin. **(A)** UMAP plot of fibroblasts subsets in snRNAseq data. **(B)** Heatmap showing expression of genes by skin fibroblast subsets. **(C)** Representative Xenium sections from each clinical category showing localization of selected fibroblast subsets. **(D-E)** Quantification of fibroblast subsets in each manually-annotated layer of skin (papillary dermis, reticular dermis, or subcutis). Statistics calculated using Kruskall-Wallis test with post-hoc Dunn testing. **(F)** Differentiation trajectories of myofibroblasts predicted by the Slingshot algorithm. **(G)** Expression of selected ISGs and fibrosis-associated genes among finely annotated subsets of reticular, inflammatory, and myofibroblasts. **(H)** Enrichment of fibrogenic and type II interferon stimulated gene (ISG) signatures in each subset of fibroblasts. *All plots:* \**p* < 0.05; \*\**p* < 0.05; \*\*\**p* < 0.005; *ns*, not significant.

Several specialized fibroblast subsets surround cutaneous blood vessels and appendages. Perivascular fibroblasts localize near endothelial cells and are distributed diffusely around the subcutaneous capillary network as well as focally around larger vessels **(Fig. 3B, C; Fig. S3B, C)**. They express lipid transport factors (*APOE, ABCA6*) and vascular basement membrane components (*COL4A2)*, with endothelial differentiation and lipid transport pathways particularly enriched in their transcriptional signature **(Fig. S3D)**. We further observed fibroblastic-reticular-cell (FRC)-like fibroblasts that share part of the perivascular FB signature and express homeostatic levels of immune-related genes like *CCL19* and *IL33* **(Fig. 3B)**. These cells localize near the superficial vascular plexus and hair follicle-associated vasculature. Interestingly, they were also enriched in TLS-like domains of diseased skin **(Fig. S3E)**. Finally, we observed Wnt-regulating fibroblasts of the dermal papilla in association with hair follicles and a population of poorly-understood *SEMA3G*+ fibroblasts in association with eccrine glands **(Fig 3B; S3D, E**), likely corresponding to recently described *CGRP*-fibroblasts (*33*). In summary, we recovered a rich diversity of homeostatic skin fibroblasts from healthy skin, each with localizations and gene expression programs tailored to their expected niche-specific functions.

### Interferon-stimulated inflammatory fibroblasts and myofibroblasts accumulate in fibrosing skin

In MSD skin, we observed a marked accumulation of two additional fibroblast subsets: inflammatory fibroblasts (iFBs) and myofibroblasts **(Fig. 3B-E)**. Inflammatory fibroblasts expressed chemokines and evidence of JAK/STAT signaling **(Fig. 3B)**, whereas myofibroblasts expressed fibroblast activation markers such as *ADAM12, LRRC15, FAP, POSTN,* similar to myofibroblasts seen in chronic inflammation and desmoplastic stroma in other organs **(Fig. 3B)** (*34*, *35*). Pathogenic fibroblast cell types were found in both inflamed and fibrotic stroma, with iFBs somewhat more enriched in inflammatory stroma and myofibroblasts in fibrotic stroma **(Fig. S3E)**.

We next compared the abundance of fibroblast populations in healthy and fibrotic skin. Pathogenic iFBs and myofibroblasts accumulated in MSD and EF samples, with this accumulation restricted to the stromal compartment affected by each disease subtype (papillary dermis in SM, reticular dermis in PM/GM, subcutis in MP/EF). For each disease subtype, the accumulation of pathogenic fibroblasts was mirrored by a loss of local FB populations at the involved stromal compartment: papillary FBs were lost in superficial morphea **(Fig. S3F)**, *PI16^+^* reticular FBs were less abundant in the reticular dermis of plaque and generalized morphea, while perivascular FBs and septal *PI16^+^* FBs were reduced in the subcutis of morphea profunda and EF skin **(Fig. 3C-E)**. We performed pseudotime trajectory analyses to gain insight into potential developmental relationships between fibroblasts. This yielded three differentiation trajectories that began in various populations of homeostatic FBs and converged on a common pathway through iFBs, terminating in myofibroblasts, with gradual acquisition of the core fibroblast activation/fibrogenesis signature **(Fig. 3F, G)**. Taken together, these data support the notion that pathogenic inflammatory fibroblasts arise from local fibroblast cell types present at the site of tissue inflammation and injury before terminally differentiating into myofibroblasts and nucleating the fibrotic niche.

We examined local inflammatory pathways that might drive homeostatic fibroblasts to adopt inflammatory and myofibroblastic identities. At the molecular level, Visium and Xenium inflammatory domains, which harbor iFBs, exhibit selective expression of canonical IFN-γ-response genes such as *CXCL9, CXCL10,* and *STAT1* **(Fig. 1H, Fig. S2F)**. Indeed, in our snRNAseq data, we found that iFBs express high levels of canonical IFN-γ-response genes and a type II interferon-stimulated gene signature **(Fig. 3G,H)**, in contrast to fibrotic fibroblasts, which are marked by fibrosis-associated genes and a fibrogenic module **(Fig. 3F,G)**.

Collectively, our analyses of skin fibroblasts in MSD suggest that IFN-γ can promote a transition from a homeostatic identity, irrespective of precise localization and homeostatic program, to an inflammatory and then myofibroblast identity.

### CD8+ T cells are significantly enriched in fibrotic skin

We next sought to determine the immunologic drivers of inflammation and fibrosis in MSD skin. Fine annotation of snRNAseq data revealed several immune cell populations that were globally increased across MSD subtypes **(Fig. S4A-D)**. A subpopulation of macrophages expressing *CCL18* was increased in lesional skin, with a reciprocal decrease in homeostatic *LYVE1*^+^ macrophages (**Fig. S4B-D)**. Although morphea and EF are not thought to be associated with significant autoantibody production, we noticed a surprising increase in antibody-secreting cells in MSD subtypes with deeper dermal and subcutaneous involvement, as well as *BCL6*^+^*CXCR5*^+^ peripheral helper T cells (Tph, **Fig. 4A**, **Fig. S4A, C, D**) (*36*). Tph cells and B cells could be found in close proximity in TLS-like aggregates, although these were present in only a subset of patients **(Fig. S4E, F)**.

**Fig. 4.**
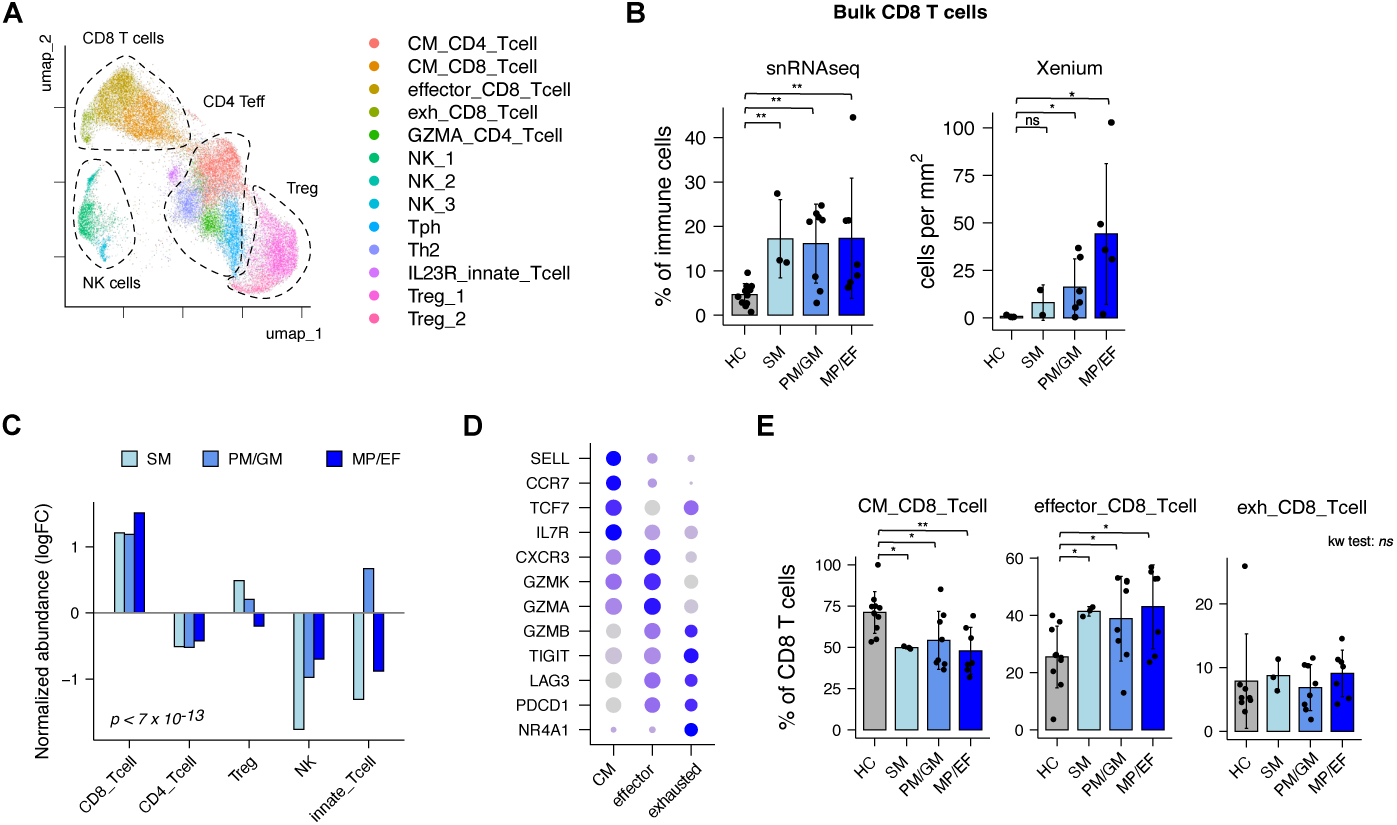
CD8 T cells are increased in fibrosing skin and orchestrate fibro-inflammatory domains. **(A)** UMAP of T cell and NK subsets in snRNAseq data. **(B)** Frequency and cell counts of bulk CD8 T cells in snRNAseq and Xenium data respectively. Statistics calculated with Kruskall-Wallis test with post-hoc Dunn testing. **(C)** Log fold change in frequency of T cell subsets as compared to healthy skin. Statistics calculated with linear mixed-effects model with disease subtype and T cell subset as fixed effects. Chi-square *p*-value for the cell type fixed effect is reported. **(D)** Expression of selected marker genes in CD8 T cell subsets. **(E)** Frequency of CD8 T cell subsets. Statistics calculated with Kruskall-Wallis test with post-hoc Dunn testing. *Inset scale bar:*20µm. *All plots:* \**p* < 0.05; \*\**p* < 0.05; \*\*\**p* < 0.005; *ns*, not significant.

The most striking and consistent change in the immune compartment of affected skin was an increase in CD8+ T cells, which increased two to four-fold across MSD disease subtypes **(Fig. 4B)**. Although bulk T cells were numerically increased in MSD skin, CD8+ T cells were disproportionately increased in frequency within the T cell compartment **(Fig. 3C)**. We found that CD8+ T cells in MSD comprised three subsets; 1) a central memory/progenitor-like phenotype marked by *TCF7*, *IL7R,* and *CCR7* expression; 2) a *GZMK*^+^ effector subset; and 3) an exhausted phenotype with higher expression of *LAG3, PDCD1,* and *NR4A1* **(Fig. 4D, Fig. S4C)** (*37*). Of these, the *GZMK*^+^ effector-like CD8+ T cells were most enriched within the CD8 compartment **(Fig. 4E)**. This subset resembled stromal-interacting CD8+ T cells seen in aging tissue and other inflammatory diseases (*38–40*), raising the possibility of direct CD8+ T cell - fibroblast interactions.

Indeed, CD8+ T cells could be seen in close proximity to disease-associated iFBs and myofibroblasts **(Fig 5A)** and were more closely co-localized with these cells than with homeostatic fibroblast populations in all MSD subtypes **(Fig. 5B, C)**. Cellchat analysis of ligand-receptor interactions predicted particularly strong interactions between myofibroblasts and CD8+ T cells, with iFB-CD8+ T cell interactions also enriched **(Fig. 5D).** CD8+ T cells were predicted to interact directly with these cells via direct production of IFN-γ, secretion of TGF-β, FASL-FAS interactions, and TRAIL-DR5 *(TNFSF10-TNFRSF10B)* (**Fig. 5E**). Given the strong upregulation of IFN-γ stimulated genes in inflammatory stroma and pathogenic fibroblasts specifically, we analyzed the IFN-γ signaling network more closely, finding that CD8+ T cells were the dominant producers of this cytokine, which signaled to numerous cell types present within inflamed stroma **(Fig. 5F).**

**Fig. 5.**
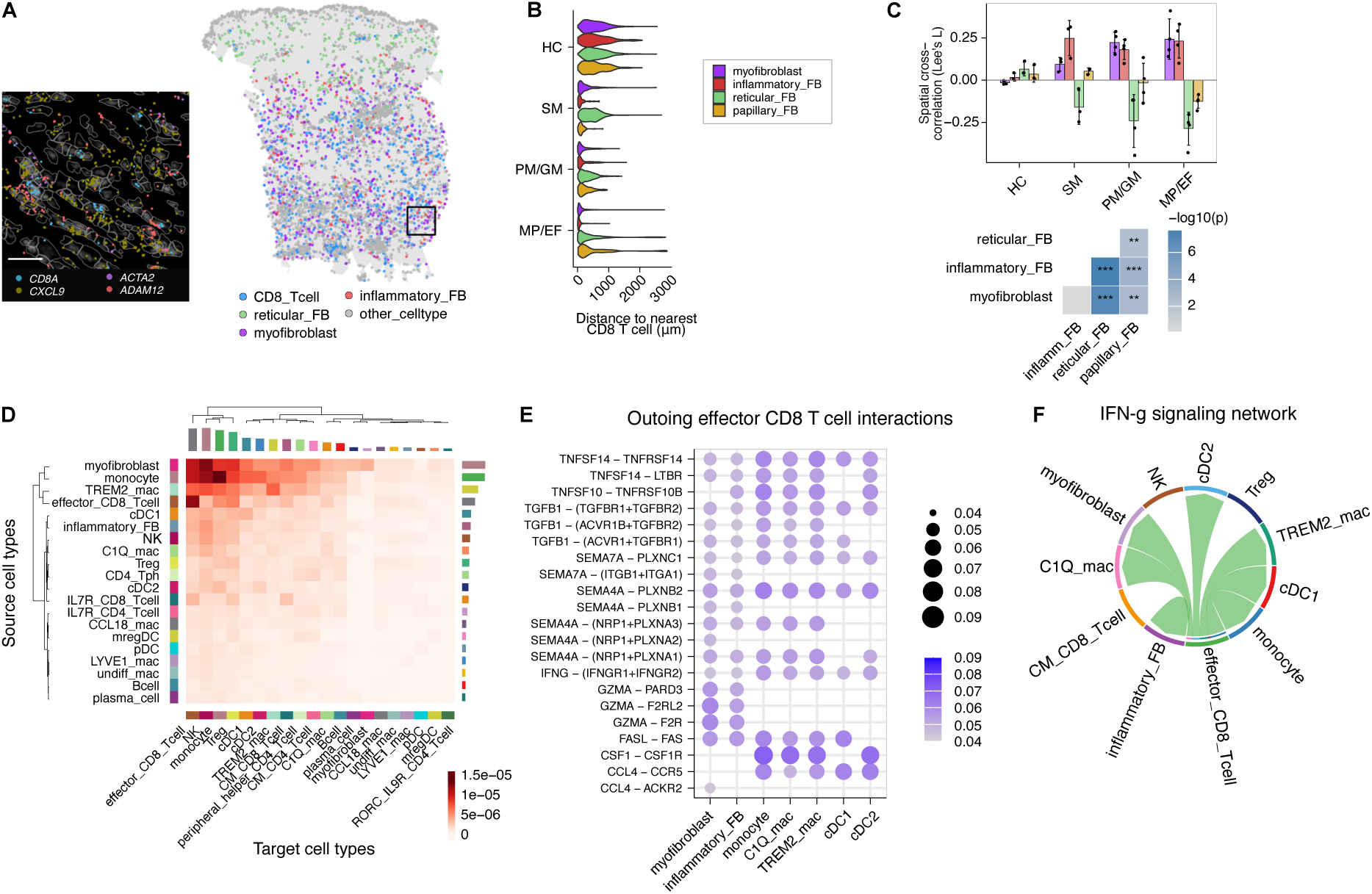
CD8 T cells co-localize and interact with pathogenic fibroblasts. **(A)** Sample Xenium image of CD8 T cell and fibroblast subset localization in eosinophilic fasciitis skin. **(B)** Distance of fibroblast subsets to the nearest CD8 T cell in Xenium data. **(C)** *Top:* Spatial cross-correlation of CD8 T cells and fibroblast subsets quantified using Lee’s *L* statistic. Statistical testing performed with a 2-way ANOVA with fibroblast cell type and disease subset as main effects (cell type effect: *p*=1.76 × 10^−13^).*Bottom:* Adjusted *p-*values of Tukey post-hoc pairwise comparisons for the cell type factor. **(D)** Predicted pairwise interaction strengths of immune and fibroblast cell types using CellChat. Cell type pairs with interaction strengths in the bottom 50^th^ percentile are not plotted. **(E)** Selected CD8 → target cell ligand-receptor interactions. **(F)** Cellchat-predicted IFN-γ signaling network. All plots: *p < 0.05; **p < 0.05; ***p < 0.005; ns, not significant.

Fibrosis has classically been associated with T_h_2 responses through the effects of IL-13 on fibroblasts. We previously found an increase in GATA3-expressing CD4+ T cells in a small cohort of patients with eosinophilic fasciitis (*41*). However, in this more robust data set, we observed that *GATA3* expression was present in numerous T cell types, a small fraction of which were bona fide T_h_2 cells expressing *IL5* and *IL13* **(Fig. S5A)**. Thus, although GATA3+ CD4+ T cells trended toward being numerically increased, true T_h_2 cells decreased as a proportion of T cells **(Fig. S5B, C)**.

### Replacement of reticular fibroblasts and enrichment of CD8 T cells in SSc skin

SSc is a devastating systemic autoimmune disease causing fibrosis of several organ systems, including the skin. SSc can present in two classic patterns of skin involvement: limited SSc (lSSc), where fibrosis is limited to the distal fingers and perioral skin, and diffuse SSc (dSSc), where fibrosis can progress proximally along limbs and involve the torso. The latter often has more severe internal organ involvement, especially interstitial lung disease, a major cause of mortality in SSc. To evaluate whether our findings apply more broadly across fibroinflammatory skin diseases, we re-analyzed a large single-cell RNA sequencing dataset from skin of 94 patients with SSc (42 lSSc, 52 dSSc) and 50 healthy donors (*42*). Our cell-type annotations differ from the original analysis, which also did not clearly distinguish T cell subsets in skin. To enable a more direct comparison between the two datasets, we performed similar processing of the SSc dataset before transferring our cell-type annotations onto skin lymphocytes and fibroblasts separately **(Fig. S6A,B, S7A-D)**. In line with the original analysis, we found broadly similar changes in fibroblast composition in SSc as in deeper forms of morphea **(Fig. S6B,C)**, with a marked reduction in *PI16+, LGR5+* reticular fibroblasts and appearance of myofibroblasts, and, to a lesser extent, iFBs **(Fig. S6C,D)**. SSc largely spares the upper dermis; accordingly, papillary fibroblasts were unaffected **(Fig. S6C)**. Furthermore, while iFBs comprised a relatively small proportion of fibroblasts in SSc patients, they strongly expressed a type 2 interferon-stimulated-gene signature **(Fig. S6E,F)**. Previous work has shown that CD8+ T cells have pro-fibrotic potential in SSc (*43–46*). In line with this work, our re-analysis of skin lymphocytes within the SSc dataset found that CD8+ T cells are significantly enriched in both lSSc and dSSc skin; as in MSD, these CD8+ T cellss strongly express *GZMK* and are the dominant cellular source of IFN-γ in skin **(Fig. S7A-D)**. Taken together, these data suggest that, in MSD as well as SSc, CD8+ T-cell-derived IFN-γ can promote a transition from homeostatic fibroblasts to iFBs and myofibroblasts.

### CD8+ T cell ablation ameliorates bleomycin-induced fibrosis in mouse skin

Our human studies detailing the major cellular and molecular constituents of skin fibrosing disease suggest that IFN-γ-secreting CD8+ T cells may play an active role in mediating the pathogenesis of these diseases. To mechanistically test the contribution of CD8+ T cells to fibrogenesis, we employed the well-established mouse model of bleomycin-induced skin fibrosis. Repetitive subcutaneous administration of bleomycin induces deep dermal and subcutaneous inflammation alongside progressive dermal fibrosis. Histopathological alterations in skin architecture are detectable as early as 5 days after treatment initiation, comprising dermal thickening and collagen deposition (*47*). To determine whether CD8+ T cells are involved in the early stages of fibrosis, we analyzed skin on day 7 of bleomycin treatment **(Fig. 6A)**. Immunohistochemical and flow cytometric analyses revealed a marked increase of CD8+ T cells within the dermis and subdermal adipose tissue compared to controls **(Fig. 6B,C; Fig. S8A,B)**. As in human skin, CD8+ T cells were major producers of IFN-γ, and bleomycin treatment enhanced this capacity **(Fig. S8C,D).** To test their functional contribution to the early fibrotic response, we depleted CD8+ T cells during bleomycin treatment **(Fig. 6D, Fig. S8A,B)**. Histological analyses of skin on day 7 demonstrated that CD8+ T cell depletion reduced dermal thickening and collagen deposition (**Fig. 6E-G**). To evaluate whether CD8+ T cell depletion induces durable changes in fibrosis, we exposed mice to bleomycin for 3 weeks. At this time point, CD8-depleted mice continued to exhibit reduced dermal thickness and collagen accumulation compared to isotype-treated controls (**Fig. 6H-K**). These data suggest that CD8+ T cells can directly contribute to fibrosis progression in skin.

**Fig. 6.**
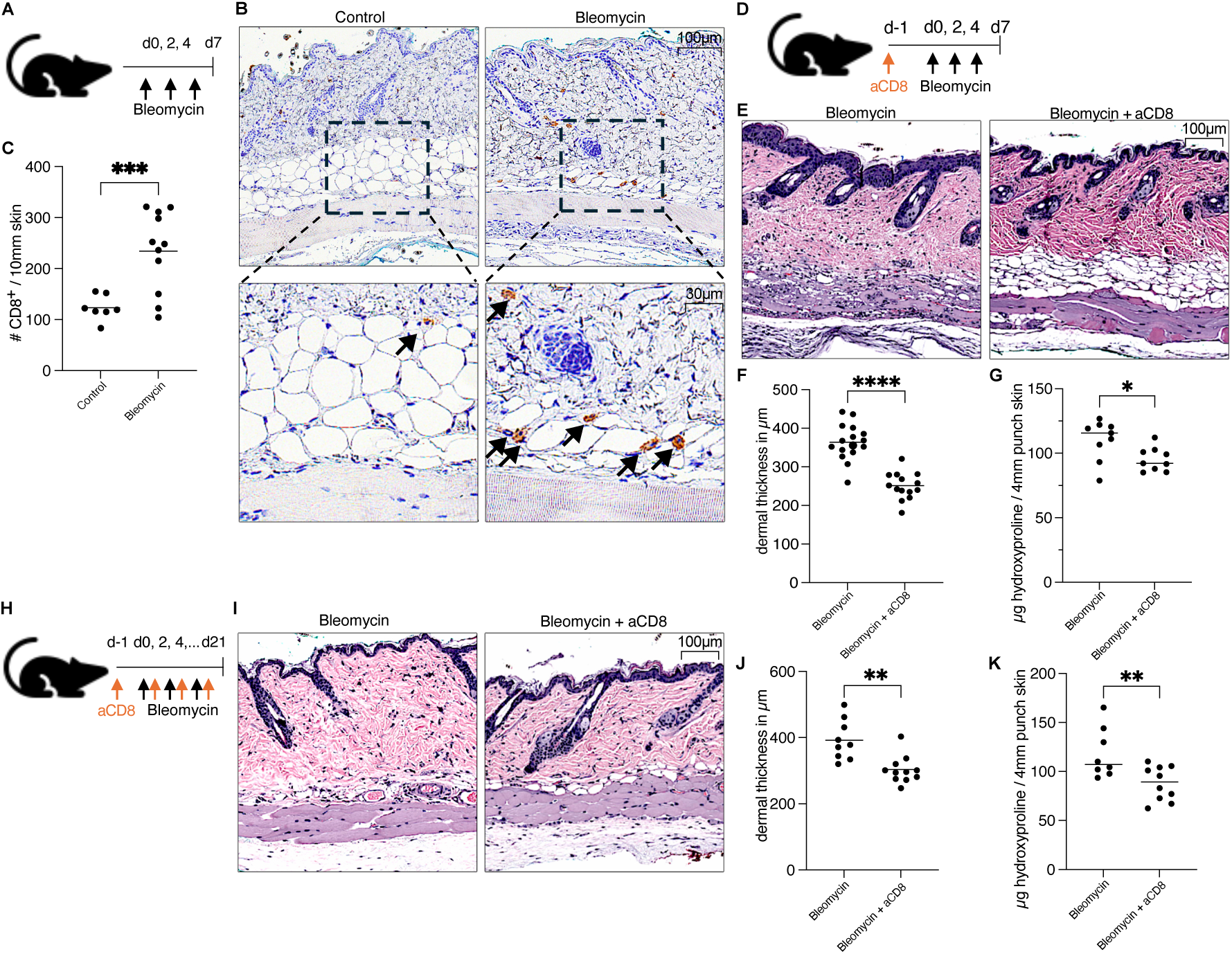
CD8 ablation ameliorates bleomycin-induced fibrosis in mouse skin. **(A)** Mice were treated with either PBS (control) or bleomycin and skin was analyzed on day 7. **(B)** Representative skin images with CD8 immunostaining and **(C)** quantification of CD8 cells in skin after PBS or bleomycin treatment. **(D-G)** Mice were treated with either bleomycin + isotype antibody or bleomycin + α-CD8 and back skin was analyzed on day 7. Representative histology (E), average dermal thickness of back skin (F) and hydroxyproline content (G) are shown. **(H-K)** Mice were treated with either bleomycin + isotype antibody or bleomycin + α-CD8 and back skin was analyzed on day 21. Representative histology (I), average dermal thickness of back skin (J), and hydroxyproline content (K) are shown. Data in C, F, G are pooled from 3 independent experiments; data in J, K are pooled from 2 independent experiments. Statistics calculated using unpaired t-tests with Welch’s correction. *All plots:* \**p* < 0.05; \*\**p* < 0.05; \*\*\**p* < 0.005; \*\*\*\**p* < 0.001; *ns*, not significant.

### IFN-γ signaling in fibroblasts promotes skin fibrosis in mice

We next sought to elucidate the mechanism by which CD8+ T cells promote fibrosis. Our prior transcriptomic analyses in human skin fibrosis identified CD8+ T cell-derived IFN-γ signaling as a driver of the inflammatory fibroblast state. We tested this in murine skin by sorting Thy-1+ PDPN+ fibroblasts from the skin of mice that were treated with bleomycin for 0, 1, 2, or 3 weeks and performing bulk RNA sequencing. In accordance with our human data, fibroblasts from bleomycin-treated skin upregulated numerous IFN-γ stimulated genes **(Fig. 7A-B)**. To directly test this pathway *in vivo*, we generated *Pdgfra^CreER/+^*, *Ifngr1^fl/fl^*mice (IFNGR-cKO), which lack IFN-γ receptor specifically in fibroblasts. IFNGR-cKO mice were treated with tamoxifen prior to subcutaneous bleomycin administration to delete IFNGR1 on the majority of fibroblasts. Skin analyses on day 7 confirmed specific depletion of the IFN-γ receptor in fibroblasts at ∼50% efficiency, without affecting immune or endothelial cell compartments **(Fig. S8E-F)**. IFNGR-cKO mice demonstrated reduced dermal thickness and collagen deposition compared to Cre-negative littermates at both the early, day 7 timepoint **(Fig. 7C-F)** and the later, day 21 timepoint **(Fig. 7G-J)**. Collectively, these data implicate a CD8+ T cell-IFN-γ-fibroblast axis in driving skin fibrosis.

**Fig. 7.**
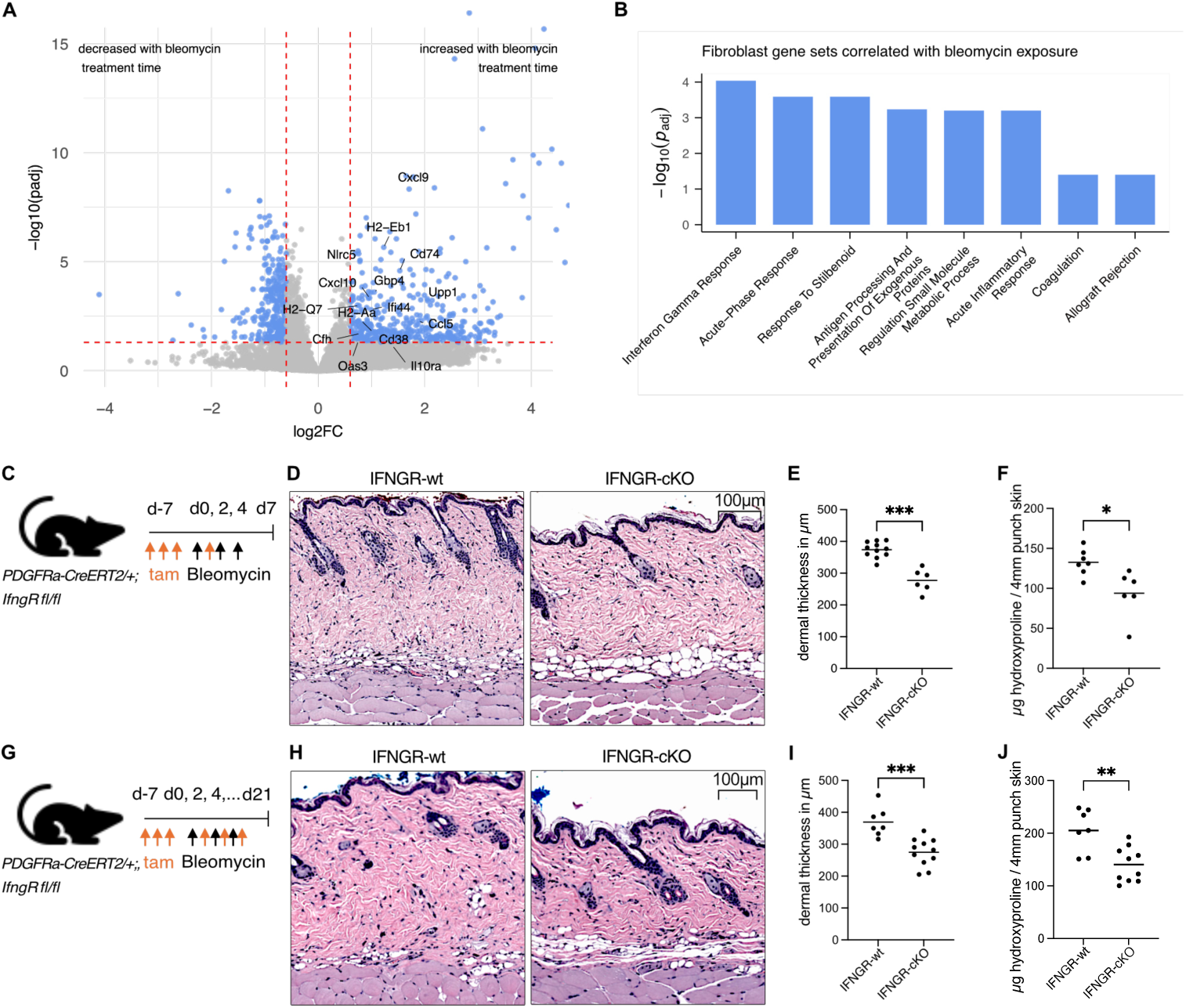
Fibroblast IFN-ɣ signaling promotes skin fibrosis in mice. **(A-B)** Mice were treated with bleomycin and fibroblasts were sorted from lesional skin at weeks 0, 1, and 2, and 3 for bulk RNA sequencing. **(A)** Differentially expressed genes associated with bleomycin treatment time. Selected IFN-γ stimulated genes are labeled. **(B)** Top GO and MsigDB gene signatures that are enriched in bleomycin-treated fibroblasts. **(C-F)** *Pdgfra^CreERT2/+^, Ifngr1^fl/fl^* and *Pdgfra^CreERT2/+-^, Ifngr1^fl/fl/f^*littermates were pre-treated with tamoxifen followed by bleomycin injections. Back skin was analyzed on day 7. Representative skin histology (D), dermal thickness (E), and hydroxyproline content (F) are shown. **(G-J)** Mice were pre-treated with tamoxifen followed by bleomycin injections as in C-F, then sacrificed on day 21. Representative skin histology (H), dermal thickness (I), and hydroxyproline content (J) are shown. Data in E, F, I, J are pooled from 2 independent experiments. Statistics calculated using unpaired t test with Welch’s correction unless indicated otherwise. \**p* < 0.05; \*\**p* < 0.05; \*\*\**p* < 0.005; \*\*\*\**p* < 0.001; *ns*, not significant.

## DISCUSSION

How the immune system initiates and sustains fibrosis remains incompletely resolved, especially across microanatomic compartments within a single organ. MSD provides an opportunity to comparatively study fibroinflammatory pathology across different skin layers.

Indeed, MSD comprises several subtypes that, while sharing core clinical, histological, and molecular features (*48*), affect the skin at different depths. Our comprehensive cross-MSD atlas revealed a consistent pattern of pathology across MSD subtypes. At the depth most affected in each MSD subtype, IFN-γ–responsive inflammatory fibroblasts and myofibroblasts replace local, homeostatic fibroblasts, with other skin layers remaining relatively unaffected. Furthermore, CD8+ T cells, the dominant cellular source of IFN-γ in lesional skin, are the most consistently and prominently expanded immune population across all disease subtypes, co-localizing with pathogenic fibroblast populations in fibroinflammatory tracts, suggesting that CD8+ T cells may act directly on fibroblasts to promote fibrotic pathology. Critically, we functionally validate these findings in mice, where CD8+ T cell depletion, as well as fibroblast-specific deletion of the IFN-γ receptor, attenuates dermal fibrosis.

As others have shown (*33*), fibroblasts in the papillary dermis, reticular dermis and subcutis differ substantially in their transcriptional programs. A possible implication of our data is that any homeostatic fibroblast population (whether in superficial, reticular, or subcutaneous compartments) may be subject to fibrotic transformation, provided that the local environment experiences the right inflammation. Our pseudotime analysis showing that, in skin, both papillary and reticular fibroblasts can serve as states of origin for iFBs, which then differentiate to myofibroblasts, buttress this idea. Recent lineage-tracing studies have described unique fibrotic fibroblast precursor populations in various tissues and models (*49–51*). Our results raise the possibility that these fibroblast precursors may give rise to myofibroblasts simply by virtue of being at the anatomic site of inflammation, rather than being uniquely poised for fibrogenesis. Combining traditional lineage tracing with spatial transcriptomics and spatial barcoding techniques may represent an avenue to definitively address this question in future studies.

Our data position CD8+ T cells as important profibrotic mediators in MSD and SSc, at least in part by acting directly on fibroblasts through IFN-γ. We do not imply that CD8+ T cells are the sole pathogenic cell type in MSD or SSc, nor that fibroblasts are their only functionally relevant target. The immune landscape of fibrosing skin is complex: we observe expansions of CCL18+ macrophages, which have known roles in fibrotic disease (*52–55*), antibody-secreting cells, and peripheral helper T cells in deeper disease subtypes. Additionally, CD8+ T cells are predicted by ligand-receptor modeling to interact with multiple cell types within fibroinflammatory zones through IFN-γ, TGF-β and death receptor signaling. While the latter two have been studied extensively in the context of fibrosis (*52*, *53*), CD8+ T cells have not attracted much attention as sources of these signals.

The dominant CD8+ T cell phenotype in lesional skin, characterized by *GZMK* expression, has also been identified in rheumatoid arthritis synovium and lupus nephritis kidneys (*39*, *56*). Intriguingly, *GZMK*+ CD8+ T cells have been described in phenotypically normal tissue in the context of aging and subclinical inflammation in mice and humans (*38*), raising the possibility that this population is not exclusively pathogenic but represents a tissue-resident sentinel that becomes dysregulated under conditions of sustained immune activation. Indeed, the origin of these cells in fibrosing skin — whether they are recruited from circulation or expand locally from tissue-resident precursors — is not resolved by the present data. Lastly, our methods did not allow us to evaluate T cell clonality. TCR sequencing and target prediction methods might determine whether distinct CD8+ T cell specificities and autoantigen localization define and perhaps drive MSD subtype.

Our *in vivo* mouse experiments strongly suggest that CD8+ T cell-derived IFN-γ plays a profibrotic role in skin in part by acting directly on fibroblasts, whereas our human results suggest the most interferon-responsive fibroblasts are iFBs. These findings may appear to conflict with a substantial prior literature demonstrating antifibrotic effects of IFN-γ *in vitro* and in some experimental models. iFBs may play two, non-exclusive fibrogenic roles: as ready substrates for myofibroblast differentiation, and as chemokine factories that recruit more fibrogenic immune cells. IFN-γ may also directly promote the myofibroblast program in some physiologic contexts, whereas supraphysiologic IFN-γ may cause distinct effects. In fact, JAK inhibitors, which inhibit IFNGR signaling, have shown clinical efficacy in patients with SSc, with selective reductions in IFN-induced gene signatures in fibroblasts and other stromal cells (*57*, *58*). We expect that elucidating precisely how IFN-γ signaling on fibroblasts promotes fibrosis will yield additional important therapeutic insights.

We acknowledge several limitations of our study. Our largely retrospective, single-nuclear and spatial methods precluded peripheral blood collection and antigen-receptor sequencing. These data might have enabled inferences regarding systemic immune activation in MSD and sources of pathogenic cell types. The mouse bleomycin model is driven by oxidative injury and DNA damage (*59*). Thus, while widely used and mechanistically informative, this model does not recapitulate the autoimmune etiology of MSD or SSc; how CD8+ T cells are recruited to skin in this model is also not clear. Nonetheless, we use this model to examine more downstream events and establish that CD8+ T cells promote fibrosis, as does IFN-γ signaling within fibroblasts.

In conclusion, this work establishes a CD8+ T cell–IFN-γ–fibroblast axis as a consistent and causally validated driver of cutaneous fibrosis across MSD and SSc.

## MATERIALS AND METHODS

### Study Design

This study aimed to determine immune and stromal drivers of skin fibrosis in MSD. We performed multi-modal transcriptomic profiling of patient skin biopsies, obtained both through pathology archives and prospective sample collection. These data were analyzed computationally to determine differences in cell abundances, transcriptional programs, cell and tissue domain localization, and cell-cell interactions between MSD subtypes and healthy controls. We then performed genetic and pharmacological manipulation of mouse models to mechanistically test hypotheses formulated from the human data.

Sample sizes for human experiments were calculated based on the number of patients needed to detect a change in population abundance between fibrosed and healthy skin at the 5% significance level with 80% power with an effect size of *f* = 1 with ANOVA testing. Some analyses, such as determination of cell and tissue domain identities and cluster abundances, were predefined. Others were selected in an exploratory fashion. Sample sizes for mouse experiments were based on prior experience with these models (*60*); dedicated power analyses were not performed again. Analyses for mouse experiments were pre-defined.

For both human and mouse study arms, data collection was not stopped early, and all predefined end points were analyzed. No data were excluded from analyses, and outliers were not removed unless they were attributable to technical failure. Numbers of human samples, animals and biological replicates for each experiment are reported in the figure legends and the Supplementary Materials. All mouse experiments were replicated at least twice. Investigators analyzing mouse experiments were blinded to group allocation.

### Human clinical samples

Skin biopsies from patients with morphea or eosinophilic fasciitis were obtained as part of a two-arm study approved by the University of California San Francisco Institutional Review Board (no. 23-38316). The first group of patients were recruited prospectively from dermatology clinic from 2024 onward, while the second group consisted of retrospectively identified tissue blocks collected between 2018 and 2024. Clinical characteristics of patients are in Supplementary Table 1. All patients met a clinical diagnosis of one or more subtypes or morphea or eosinophilic fasciitis with active lesions at the time of biopsy. Patients receiving systemic biologic therapy (*e.g.* intravenous immunoglobulin or dupilumab), targeted synthetic oral therapy (*e.g.* JAK inhibitors), or oral steroids in excess of 0.5m/kg/day were excluded. All specimens were reviewed by a board-certified dermatopathologist (JNC) to confirm concordant histopathology. Healthy control tissue was obtained from surgical discard specimens.

### Mice

C57BL6/J (Wild-type) were purchased from Jackson Laboratories. B6.129S-Pdgfratm1.1(cre/ERT2)Blh/J (*Pdgfra^CreERT2^*), and C57BL/6N-Ifngr1tm1.1Rds/J (*Ifngr1^flox^*) mice were a gift from Ari Molofsky. All mice were bred and maintained in a specific-pathogen-free mouse facility in accordance with an animal protocol approved by the Institutional Animal Care and Use Committee of the University of California, San Francisco (protocol AN204200). Mice were socially housed under a 12-h day/night cycle at 25 °C and ambient humidity. Littermates were used as controls, and animals of both sexes were included. Parental cage and weaning cage were randomized among experimental groups.

### Single-nuclear RNA sequencing, preprocessing, and annotation

Paraffin-embedded biopsies were sectioned into 30µm thick tissue scrolls, and three scrolls per specimen were collected in Miltenyi GentleMACS C-tubes. Nuclei were extracted using 10X Genomics protocol CG000632. Briefly, tissue scrolls were de-paraffinized with xylene and ethanol, and tissue was digested using Liberase TH (Sigma) on a GentleMACS OctoDissociator. Nuclei were passed through a 30µm filter and counted on a Cellaca MX counter. Probe hybridization and library preparation performed using the 10X Genomics Chromium Fixed RNA Profiling kit. An average of 18,000 nuclei per sample were pooled in each library. One 4-plex library and two 16-plex libraries were sequenced on an Illumina NovaSeq X, with a median of 11,493 nuclei recovered per sample.

Fastq files were then aligned to human genome assembly GRCh38 and demultiplexed using Cellranger 8.0.0. Cellbender 0.3.0 was used to remove ambient RNA and call empty droplets, using parameters --expected-cells of 600 and --total-droplets-included of 17000. Barcode-count matrices were loaded into Seurat 5.2.1. Nuclei with fewer than 750 reads and/or mitochondrial content between 4 - 9% were discarded. Same-sample multiplets were removed using DoubletFinder. Data were normalized and scaled using the NormalizeData, FindVariableFeatures, and ScaleData functions, then integrated using the harmony package with patient ID as the grouping variable. The FindClusters function was used to perform clustering. Coarse clustering was initially performed to identify immune, stromal vascular, and epidermal subsets, which were then split into separate Seurat objects. Cell subsets were iteratively identified, re-clustered, and manually annotated to yield fine cell annotations (Immune object → T cell, macrophage, B cell objects; stromal object → fibroblast object). At each sub-clustering stage, doublets were manually identified and removed, and superset objects (immune, stromal) were re-constituted by merging the constituent sub-objects.

To score fibrosis and IFN-γ signatures, we used the Seurat AddModuleScore function to calculate enrichment of a published fibrosis gene set (*61*) and the MsigDB Hallmark Interferon Gamma gene set, respectively.

### Pseudotime trajectory analysis

Seurat data objects containing fibroblasts identified from snRNAseq data were converted to SingleCellExperiment format. Slingshot 2.10 was run using UMAP dimensional reduction coordinates and myofibroblasts as the start cluster.

### Visium spatial transcriptomics, preprocessing, and annotation

Paraffin-embedded biopsies were sectioned at 5µm thickness onto SuperFrost slides, stained with hematoxylin and eosin, and mounted with coverslips by the UCSF dermatopathology.

These slides were first imaged on a Leica Versa DM6 slide scanner and then prepared according to the standard 10X Genomics Visium CytAssist v2 protocol (CG000518, CG000495). Briefly, tissue sections were de-coverslipped, de-paraffinized in xylene and ethanol, de-stained, de-crosslinked, and transferred to 6.5mm Visium slides with 55µm spot size. Probe hybridization, ligation, and library preparation was then performed per protocol, and libraries were sequenced on an Illumina NovaSeq X.

Fastq files were aligned using SpaceRanger 2.0.1. Due to poor automated detection of subcutaneous tissue, we performed manual tissue detection in the 10X Loupe Browser prior to running SpaceRanger. Spot-count matrices were loaded into Seurat 5.2.1 and filtered with a count threshold of 500. The data were then processed in a similar fashion to snRNAseq data with manual cluster annotation.

### Visium cell type frequency prediction

The proportions of cell types within each Visium spot were estimated using the RCTD algorithm (*62*). snRNAseq data was downsampled to 10,000 cells while preserving cell type proportions; this was used as a reference dataset for cell type deconvolution. RCTD was then run using default parameters with doublet_mode set to “full.”

### Visium spatial co-localization analysis

We used Lee’s *L* statistic, a measure of bivariate spatial association, to quantify the spatial association of cell types in Visium data. For each Visium image, a list of nearest the immediate neighbors of each Visium spot was calculated with spdep::dnearneigh, and then used to construct a spatial weights graph with spdep::nb2listw. Each spot was also appended to its neighbor list and given a weight of 2, such that cell type colocalization within the same spot would be appropriately counted. The spatial weights graph was then used to compute Lee’s *L* statistic then for each pair of cell types using the spdep::lee function.

### Visium pseudobulk differential gene expression

Visium spots were grouped by tissue domain and mean raw UMI counts were calculated for each sample. Differential expression testing of this pseudobulked data was carried out using DESeq2 with the formula ∼ patient_id + domain. *P-*values were calculated using Wald testing and adjusted with Benjamini-Hochberg corrections.

### Xenium segmentation, clustering, annotation, and domain detection

FFPE skin biopsies were sectioned at 5µm thickness and deposited onto Xenium slides (3 biopsies per capture area). Slides were then prepared for spatial transcriptomics according to the standard 10X Genomics Xenium In Situ protocol (CG000749). Briefly, sections were de-paraffinized, permeabilized, and hybridized with a custom 480-gene Xenium v1 panel (**Supplementary Table S2**). Following probe ligation and signal amplification, the Xenium Multi-Tissue Cell Segmentation Kit was applied. Slides were then imaged on the Xenium Analyzer (CG000584). Onboard analysis performed using Xenium Onboard Analysis (XOA) software, using nuclei-, boundary-, and interior-based cell segmentation.

Downstream analysis was performed with Seurat 5.2.1. Cells with <20 transcripts or area <10µm2 were removed. Cells lying outside the main tissue area were identified and removed by computing the k-nearest neighbor distance for each cell centroid and flagging cells whose kNN distance exceeded 1.5 times the interquartile range (90th minus 10th percentile). Data were then preprocessed, integrated, and manually annotated in a similar manner to snRNAseq data. Tissue domains were identified using the Tessera package (*32*). Tessera tiles were constructed from harmony embeddings using prune_min_cells = 3, prune_thresh_quantile = 0.95, alpha = 2, min_npts = 5, and max_npts = 25. Tiles were then clustered using the Seurat FindClusters function and manually annotated.

### snRNAseq - Xenium label transfer

Diffusion-based label transfer was used to transfer snRNAseq annotations to Xenium cells using a method adapted from Stein et al (*32*). T cells, myeloid cells, B cells, fibroblasts, and non-fibroblast stromal cells were subsetted and individually subjected to label transfer. For each subset, Xenium and snRNAseq data were harmony-integrated using the 480 genes in the Xenium panel. A UMAP graph was constructed using the uwot package, and a decay factor of 0.8 was applied to the resulting adjacency matrix to yield a diffusion matrix !. Cell type label probabilities were then propagated from snRNAseq data to Xenium data using an iterative diffusion process. The cell-label matrix of the multimodal integrated object was used as the initial matrix “^!^. At each step #, the cell-label matrix “^“^was re-injected with the initial matrix and multiplied by the adjacency matrix such that “^“^ = ! ∗ (“^(“$%)^ + “^!^). After 20 iterations, transferred annotations were extracted from the final cell-label matrix and added to the metadata of the original Xenium object.

### Xenium gene imputation

Imputation of unmeasured genes in the Xenium data was then conducted using the CytoSPACE package (*63*). Multimodal annotations as calculated by label transfer above were used to guide assignment of single cells in the snRNASeq dataset to spatial locations in the Visium dataset.

CytoSPACE was run on a sample-by-sample basis to preserve biological variability. This allowed the Xenium feature list to be expanded from 480 measured genes to 480 measured genes + 17,594 imputed genes. The resulting imputed data was loaded into Seurat, preprocessed, and clustered at a resolution of 0.4. A small proportion of imputed cells did not cluster near major cell clusters in UMAP space. These poorly-clustered cells were programmatically isolated from major clusters and grouped into low-count clusters as follows. First, the shared nearest neighbors graph from UMAP coordinates was recalculated with reduced k.param of 12 and higher prune.SNN of 0.12 to remove weak edges. Next, an elbow plot of cluster size versus cumulative percentage of total cells was constructed. This clearly demonstrated an “elbow” of low-count clusters with a cumulative cell count of <5% of all cells; clusters below this threshold were filtered from the data.

### Spatially-informed cell-cell interaction analysis

Xenium data with missing genes imputed by CytoSPACE was used for CellChat analysis (*64*). Only cells within the inflamed stromal tissue domain of morphea and EF samples were included in the analysis to enrich for discovery of interactions between pathogenic immune and stromal cell subtypes, because these cells were not abundant in most other tissue domains or in healthy controls. truncatedMean was used to summarize gene expression. Cellchat communication probabilities were calculated using an interaction range of 250µm, a contact range of 20µm for juxtacrine/paracrine interactions, a spot size of 10µm, scale.distance of 1.5, and a conversion_factor of 1. Interactions with fewer than 10 cells were removed.

### Tissue domain enrichment analysis

Enrichment scores were used to quantify the observed frequency of cell types in each tissue domain compared to the frequencies expected by chance, using a method from Stein et al (*32*). Expected frequencies for cell type *i* in domain *j* were calculated as 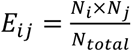 for population cell counts *N*. Enrichment scores were then calculated as 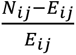; absolute values of enrichment scores were log-transformed for heatmap plotting.

### Re-analysis of published systemic sclerosis data

We downloaded the raw matrix from GEO (GSE195452) along with the cell level annotations and metadata. The raw dataset was subsetted to cells in the provided annotations, and cells with outlier values of high nCount_RNA (>15,000) and nFeature_RNA (>5,000) were removed. Following this, we processed the data using the standard Seurat (v4.3.0) workflow; we log-normalized, identified variable features, and scaled the data (additionally regressed out nFeature and nCount_RNA as well as percent.mt, percent.ribo, and cell cycle G2M and S scores) prior to running PCA. We batch-corrected the PCs using Harmony (v1.2.0) with SeqBatchID as the batch variable. A UMAP embedding was generated from the first 30 Harmony adjusted-PCs, followed by a neighborhood graph construction, and Leiden clustering (with method=’igraph’).

Annotations were summarized to major coarse populations and the data subset to SSc and HC skin samples only. One HC individual was removed because they had less than 50 cells. The object was then divided into CD45 and CD90 fractions. SingleR (v2.2.0) was then used to perform a label transfer from the annotations (separated into immune or stromal) in this paper to the appropriate Gur et al. objects. The CD90 fraction was then subset to cells annotated as fibroblasts by Gur et al.

For the fibroblast analysis, CD90 cells were filtered to those annotated as fibroblasts in label transfer and to individuals with more than 30 cells. This resulted in 51,194 cells across 139 individuals (53 HC, 38 lSSc, 48 dSSc). Fibroblast subtype proportions calculated based on the resulting cells. The IFN-γ module score was calculated in a similar fashion to the morphea/EF dataset described above.

For the T cell analysis, the lymphocyte compartment was further isolated by retaining cells annotated as T cells, B cells, NK cells, and plasma cells. Each subset underwent re-normalization and dimensionality reduction (UMAP via Harmony batch correction). Cell type labels were transferred to the lymphocyte subset using SingleR with reference-based label assignment; low-confidence calls were pruned based on delta-score thresholds. CD8+ T cell abundance was quantified per patient as the fraction of total lymphocytes, restricting to patients with >30 recovered lymphocytes (n = 81: 25 HC, 23 lSSc, 33 dSSc). CD8+ T cell fractions were compared across disease groups (HC, lSSc, dSSc) by Wilcoxon test.

### Bleomycin-induced fibrosis and injection of pharmacological agents

Back skin of mice 7-10 weeks of age was shaved, and 0.1U bleomycin sulfate (Northstar Rx) in 100 uL DPBS (ThermoFisher) was injected intradermally into a 1×1cm patch of skin three times a week on non-consecutive days for indicated length of treatment, using different needle entry points for consecutive injections. Control mice received PBS. For CD8_+ T cell depletion, mice received weekly IP injections of anti-CD8α monoclonal antibody (Bio-Xcell) at 16ug/g bodyweight, starting one day prior to the bleomycin treatment. Control mice received an isotype control (Bio-Xcell). For induction of CreER-mediated recombination, mice received tamoxifen (Sigma) dissolved in corn oil (Sigma) at 100mg/kg intraperitoneally for 3-5 days, as indicated.

### Murine skin imaging quantification

For hematoxylin and eosin (H&E) staining and immunohistochemistry, skin samples were fixed for 24-48H in 4% paraformaldehyde. H&E staining was performed through UCSF Dermatopathology Lab using standard methods, and slides were imaged on a Leica Versa slide scanner. Mouse CD8 immunohistochemistry was performed through HistoWiz (NY) using Cell Signaling Technologies rabbit anti-CD8α (clone D4W2Z). Quantification was performed using QuPath 0.5.1. For dermal thickness, distance from dermal-epidermal junction to subcutis (dermal white adipose tissue, or, when completely absent due to bleomycin, panniculus carnosus) was measured; at least 5 measurements per slice were averaged. For adipocyte quantification, an ImageJ script was implemented to detect and annotate adipocytes within a set length of the skin. White areas with area <100um^2^ were excluded. For CD8+ cell quantification, positive cells were detected and annotated using the ‘Positive cell detection’ command of QuPath. Both adipocyte and CD8+ cell annotations were then counted and normalized to a skin length of 10mm.

### Hydroxyproline assay

Hydroxyproline assays were performed as previously described (*65*). Briefly, 4mm skin biopsies were hydrolyzed in 200 uL 6N HCl at 95C overnight, then diluted 1:20 in 6N HCl again. 10µL of standards or diluted hydrolyzed samples were pipetted in duplicate into a 96-well optically clear plate with 30 μl of citric acid buffer; 100 uL of Chloramine T solution was added. After 20 min of oxidation at room temperature, 100µL of Ehrlich’s Reagent was added and plate incubated at 65°C for 20 min. Absorbance was read at 550 nm in a spectrophotometer (Beckman), and samples were quantified according to the standard curve.

### Mouse skin single cell suspension preparation and flow cytometry

For preparation of single-cell suspensions from bleomycin lesions, lesions (∼1.25×1.25cm squares of treated back skin) were dissected, weighed, and placed in 1mL of digestion medium (*41*) in 24 well plates on ice. In some experiments, subcutis was separated from the dermal/epidermal fraction, as previously described (*41*). After all tissues were harvested, skin was finely minced in the well plate, then incubated at 37C for 60min in a humidified tissue culture incubator with 95% O_2_/5% CO_2_, then placed back on ice. Resulting suspensions were filtered into 50 mL conicals through 40um mesh filters, which were then washed with 5mL cold DPBS. Suspensions were then centrifuged at 400g for 10 min. For experiments involving intracellular cytokine staining, suspensions were plated in a 96-well round bottom plate, resuspended in 1× cell stimulation cocktail (Tonbo Biosciences) in C10 medium, and incubated at 37 °C for 2.5 h before antibody staining. For antibody staining, cells were resuspended in 2% FBS in DPBS + anti-mouse CD16/CD32 (clone 2.4G2, 50ug/mL, BioXcell) and incubated on ice for 15 min. Surface antibody cocktail was prepared as 2x concentrate in 2% FBS in PBS, added to samples in anti-CD16/32 mix; samples were then incubated for 20 min on ice. For biotinylation steps, cells were washed of surface antibody cocktail twice, then incubated in streptavidin-Alexa647 (Jackson Immunoresearch, 2.5ug/mL in 2% FBS in PBS) for 15 min on ice. Live/dead staining was performed with DAPI (Tocris) or Ghost violet 510 (Fisher) as indicated. For intracellular target staining, cells were then fixed and permeabilized using a Cytofix/Cytoperm kit (BD Biosciences, 554714). Samples were run on a BD Fortessa cytometer.

The following antibodies were used: anti-CD45 Alexa700 (clone 30-F11, 1:200, eBiosciences), anti-CD31 BV711 (clone 390, 1:500, BD), anti-CD90.2 PerCP-Cy5.5 (clone 53-2.1, 1:500, Biolegend), anti-CD90.2 APC (clone 53-2.1, 1:500, eBiosciences), anti-CD90.2 eF450 (clone 53-2.1, 1:1000, Biolegend), anti-CD140a BV650 (clone APA5, 1:200, BD), anti-PDPN PE-Cy7 (clone ebio8.1.1, 1:400, Invitrogen), anti-CD3e FITC (clone 500A2, 1:200), anti-CD4 BV605 (clone RM4-5, 1:400, Biolegend), anti-CD4 BV650 (clone RM4-5, 1:400, Biolegend), anti-CD8a Alexa594 (clone 53-6.7, 1:400, Biolegend), anti-CD8a BV785 (clone 53-6.7, 1:300, Biolegend) anti-IFNGR1 biotin (clone GR20, 1:400, BD), anti-IFNg PE (clone XMG1.2, 1:100, eBiosciences), anti-TCRβ PerCP-Cy5.5 (clone 1A8, 1:400, Biolegend), anti-CD3ε BV711 (clone 145-2C11, 1:200, BD), anti-CD11b APC-eFluor780 (clone M1/70, 1:500, eBiosciences). Cell numbers were quantified using CountBright counting beads (ThermoFisher). Flow cytometry analyses were performed using FlowJo v10.10 or FlowJo v11.1.

### Bulk RNA sequencing of murine fibroblasts

Mice were subjected to skin bleomycin model, staggering the start of bleomycin injections such that, for each cohort, mice for all time points (0, 1, 2, 3 weeks of bleomycin treatment) were processed in parallel. The subcutis was manually dissected from the dermis and single-cell suspensions were prepared from each fraction of lesional skin as detailed above. Fibroblasts were sorted as live (DAPI-), CD45-, CD31-, Thy1+, PDPN+ cells using an 85um nozzle on a FACSAria III sorter (BD) directly into 6µL of 1x complete lysis buffer from low-input mRNA-seq kit (Takara), with 200-750 cells per tube. Cells were flash frozen on dry ice and stored at −80C until all cohorts were processed. For library preparation, tubes were thawed on ice, volume adjusted to 12.5uL as required, first-strand synthesis and initial PCR amplification was performed according to manufacturer’s instructions, using 14 cycles of amplification as empirically determined through pilot experiments (data not shown). Barcoded sequencing adapters (Illumina) were ligated to PCR fragments, and libraries were further amplified and quantified using Nextera XT reagents (Illumina). Final libraries were sequenced through Novogene (Sacramento, CA, USA) on a single lane of a Novaseq X 10B flow cell using paired-end 150bp protocol.

### Mouse bulk RNA sequencing analysis

Fastq files were pseudo-aligned to mouse genome assembly GRCm39, Ensembl release 110, using Kallisto. Transcript-level abundances were imported and summarized to the gene level using tximport, with counts scaled by transcript length (lengthScaledTPM). Lowly expressed genes were excluded prior to downstream analysis, retaining only those with counts per million (CPM) exceeding 1 in at least 5 samples. Counts matrices were imported to DESeq2 and analyzed with the formula ∼ timepoint * celltype, with timepoint as a continuous variable and celltype as a categorical variable (dermal or subcutaneous fibroblast). Differentially expressed genes in Figure 5 were identified by extracting the timepoint coefficient from the model using results(dds, name = ‘timepoint’), which models the change in gene expression per unit time of bleomycin treatment.

## List of supplementary materials

Fig S1 to S8

Table S1-S2

## ACKNOWLEDGEMENTS

We thank Ari Molofsky for helpful comments on the manuscript.

We thank Jamie Chan, Dacia Miyake-Caballero, Russell Lam from the UCSF Dermatopathology Service for support in acquiring human samples and mouse histology.

We thank Andrea Barczak from the UCSF Bakar ImmunoX Initiative for guidance and support in the funding from that entity.

UCSF Flow Cytometry Colab supported in part by NIH grant P30 DK063720 and by the NIH instrumentation grant 1S10OD021822-01.

Sequencing was performed at the UCSF Center for Advanced Technology, supported by UCSF PBBR, RRP IMIA, and NIH 1S10OD028511-01 grants.”

AJC is supported by NIH R35 CA24244, R01 HL181372, R01 HL175312, R35 HL171251, and support from the Melanoma Research Alliance, the UCSF Program for Breakthrough Biomedical Research, The Leo Foundation, The Cancer Research Institute, and the UCSF Bakar ImmunoX Initiative.

## FUNDING

American Partnership for Eosinophilic Diseases Hope Pilot Grant (MJK, MDR)

Bakar ImmunoX CoPilot Grant (MJK, MDR)

Mt Zion Health Fund Pilot Grant 20231627 (MJK)

National Institutes of Health P30AR070155 Pilot Award (MJK)

National Institutes of Health 5K08AR085193 (MJK)

Rheumatology Research Foundation Scientist Development Award (MJK)

Deutsche Forschungsgemeinschaft 538205505 (TCG)

National Institutes of Health 1R01AR077553 (MDR)

## AUTHOR CONTRIBUTIONS

MJK, ICB, AH, and MDR conceived and supervised the project. ICB, MJK, and AJC designed human experiments. ICB, MJK, JNC, IN, and AH conducted patient enrollment and retrospective specimen identification and retrieval. ICB, VJ, SS, and LM executed human specimen processing and data acquisition with supervision from AJC and WE. Human data analysis was performed by ICB. Analysis of published SSc data was performed by EF. TCG and MJK designed, executed, and analyzed the mouse experiments. SY and GKF provided data curation support. ICB, MJK, and MDR secured funding. MJK, ICB, and TCG wrote the manuscript draft. All authors reviewed and edited the manuscript.

## COMPETING INTERESTS

AJC has received consulting fees from Survey Genomics.

MJK is a consultant and cofounder of Radera Bio Inc.

MDR is a consultant and cofounder of TRex Bio Inc., Sitryx Bio Inc., and Radera Bio Inc. He is also a consultant for Mozart Bio Inc.

JNC is a consultant and cofounder of Radera Bio Inc and a consultant for TRex Bio Inc. All other authors declare they have no competing interests.

## DATA AND MATERIALS AVAILABILITY

All raw fastq sequencing files and Xenium Analyzer output files will be made available at the NIH database of genomes and phenotypes (dbGAP) repository prior to final manuscript publication. Previously published data used in this study can be found at NCBI Gene Expression Omnibus (accession number GSE195452). All mouse alleles used are available from the Jackson Laboratories. Mouse data will be available upon request to corresponding authors.

## REFERENCES

1. N. Fett, V. P. Werth, Update on morphea: Part I. Epidemiology, clinical presentation, and pathogenesis. Journal of the American Academy of Dermatology 64, 217–228 (2011).

2. M. Huynh, E. Bogdanski, T. Fleshman, J. Schneider, K. Libson, V. P. Werth, J. Lin, A. M. Korman, Eosinophilic Fasciitis: New Developments and Future Directions. International Journal of Dermatology 64, 1356–1370 (2025).

3. C. Papara, D. A. De Luca, K. Bieber, A. Vorobyev, R. J. Ludwig, Morphea: The 2023 update. Front. Med. 10 (2023).

4. J. Varga, D. Abraham, Systemic sclerosis: a prototypic multisystem fibrotic disorder. J. Clin. Invest. 117, 557–567 (2007).

5. T. Stein, P. Cieplewicz-Guźla, K. Iżykowska, M. Pieniawska, R. Żaba, A. Dańczak-Pazdrowska, A. Polańska, What Is New in Morphea—Narrative Review on Molecular Aspects and New Targeted Therapies. Journal of Clinical Medicine 13, 7134 (2024).

6. T. A. Wynn, Fibrotic disease and the TH1/TH2 paradigm. Nat Rev Immunol 4, 583–594 (2004).

7. S. A. Jimenez, B. Freundlich, J. Rosenbloom, Selective inhibition of human diploid fibroblast collagen synthesis by interferons. J. Clin. Invest. 74, 1112–1116 (1984).

8. M. l. Stephenson, S. m. Krane, E. p. Amento, P. a. McCroskery, M. Byrne, Immune interferon inhibits collagen synthesis by rheumatoid synovial cells associated wth decreaded levels of the procollagen mRNAs(FEBS 2156). FEBS Letters 180, 43–50 (1985).

9. E. P. Amento, A. K. Bhan, K. G. McCullagh, S. M. Krane, Influences of gamma interferon on synovial fibroblast-like cells. Ia induction and inhibition of collagen synthesis. (1985). 10.1172/JCI112041.

10. M. R. Duncan, B. Berman, Gamma interferon is the lymphokine and beta interferon the monokine responsible for inhibition of fibroblast collagen production and late but not early fibroblast proliferation. J Exp Med 162, 516–527 (1985).

11. J. Rosenbloom, G. Feldman, B. Freundlich, S. A. Jimenez, Inhibition of excessive scleroderma fibroblast collagen production by recombinant γ-interferon: Association with a coordinate decrease in types I and III procollagen messenger RNA levels. Arthritis & Rheumatism 29, 851–856 (1986).

12. J. G. Clark, T. F. Dedon, E. A. Wayner, W. G. Carter, Effects of interferon-gamma on expression of cell surface receptors for collagen and deposition of newly synthesized collagen by cultured human lung fibroblasts. J Clin Invest 83, 1505–1511 (1989).

13. J. Varga, A. Olsen, J. Herhal, G. Constantine, J. Rosenbloom, S. A. Jimenez, Interferon-γ reverses the stimulation of collagen but not fibronectin gene expression by transforming growth factor-β in normal human fibroblasts. European Journal of Clinical Investigation 20, 487–493 (1990).

14. A. S. Narayanan, J. Whithey, A. Souza, G. Raghu, Effect of gamma-interferon on collagen synthesis by normal and fibrotic human lung fibroblasts, Chest. 101 (1992)pp. 1326–31.

15. S. N. Giri, D. M. Hyde, Jr. Marafino B. J., Ameliorating effect of murine interferon gamma on bleomycin-induced lung collagen fibrosis in mice, Biochem Med Metab Biol. 36 (1986)pp. 194–7.

16. R. D. Granstein, G. F. Murphy, R. J. Margolis, M. H. Byrne, E. P. Amento, Gamma-interferon inhibits collagen synthesis in vivo in the mouse. J. Clin. Invest. 79, 1254–1258 (1987).

17. D. M. Hyde, T. S. Henderson, S. N. Giri, N. K. Tyler, M. Y. Stovall, Effect of Murine Gamma Interferon on the Cellular Responses to Bleomycin in Mice. Experimental Lung Research 14, 687–704 (1988).

18. G. Gurujeyalakshmi, S. N. Giri, Molecular Mechanisms of Antifibrotic Effect of Interferon Gamma in Bleomycin-Mouse Model of Lung Fibrosis: Downregulation of TGF-β and Procollagen I and III Gene Expression. Experimental Lung Research 21, 791–808 (1995).

19. S. D. Oldroyd, G. L. Thomas, G. Gabbiani, A. M. El Nahas, Interferon-γ inhibits experimental renal fibrosis. Kidney International 56, 2116–2127 (1999).

20. H. W. Chen, J. L. Zhu, V. Martyanov, L. C. Tsoi, M. E. Johnson, G. Barber, D. Popovich, J. C. O’Brien, J. Coias, N. Cyrus, N. Malviya, S. Florez-Pollack, E. Kunzler, G. A. Hosler, J. E. Gudjonsson, D. Khanna, M. Whitfield, H. T. Jacobe, Gene Expression Signatures in Inflammatory and Sclerotic Morphea Skin and Sera Distinguish Morphea from Systemic Sclerosis. Journal of Investigative Dermatology 143, 1886–1895.e10 (2023).

21. B. Freundlich, S. A. Jimenez, V. D. Steen, T. A. Medsger, M. Szkolnicki, H. S. Jaffe, Treatment of systemic sclerosis with recombinant interferon-gamma. A phase I/II clinical trial. Arthritis Rheum 35, 1134–1142 (1992).

22. P. G. Vlachoyiannopoulos, N. Tsifetaki, I. Dimitriou, D. Galaris, S. A. Papiris, H. M. Moutsopoulos, Safety and efficacy of recombinant gamma interferon in the treatment of systemic sclerosis. Annals of the Rheumatic Diseases 55, 761–768 (1996).

23. R. P. Polisson, G. S. Gilkeson, E. H. Pyun, D. S. Pisetsky, E. A. Smith, L. S. Simon, A multicenter trial of recombinant human interferon gamma in patients with systemic sclerosis: effects on cutaneous fibrosis and interleukin 2 receptor levels. J Rheumatol 23, 654–658 (1996).

24. N. Hunzelmann, S. Anders, G. Fierlbeck, M. Albrecht, S. Bell, J. Thur, B. Adelmann-Grill, Multicenter Trial of 1 Year of Treatment With Recombinant Interferon Gamma. doi: 10.1001/archderm.133.5.609.

25. N. Hunzelmann, S. Anders, G. Fierlbeck, R. Hein, K. Herrmalm, M. Albrecht, S. Bell, R. Muche, J. Wehner-Caroli, W. Gaus, T. Krieg, Double-blind, placebo-controlled study of intralesional interferon gamma for the treatment of localized scleroderma. Journal of the American Academy of Dermatology 36, 433–435 (1997).

26. A. Grassegger, G. Schuler, G. Hessenberger, B. Walder-Hantich, J. Jabkowski, W. MacHeiner, W. Salmhofer, B. Zahel, G. Pinter, M. Herold, G. Klein, P. O. Fritsch, Interferon-gamma in the treatment of systemic sclerosis: a randomized controlled multicentre trial, Br J Dermatol. 139 (1998)pp. 639–48.

27. Jr. King T. E., C. Albera, W. Z. Bradford, U. Costabel, P. Hormel, L. Lancaster, P. W. Noble, S. A. Sahn, J. Szwarcberg, M. Thomeer, D. Valeyre, R. M. du Bois, Effect of interferon gamma-1b on survival in patients with idiopathic pulmonary fibrosis (INSPIRE): a multicentre, randomised, placebo-controlled trial, Lancet. 374 (2009)pp. 222–8.

28. W. E. Thierfelder, J. M. van Deursen, K. Yamamoto, R. A. Tripp, S. R. Sarawar, R. T. Carson, M. Y. Sangster, D. A. A. Vignali, P. C. Doherty, G. C. Grosveld, J. N. Ihle, Requirement for Stat4 in interleukin-12-mediated responses of natural killer and T cells. Nature 382, 171–174 (1996).

29. M. H. Kaplan, Y.-L. Sun, T. Hoey, M. J. Grusby, Impaired IL-12 responses and enhanced development of Th2 cells in Stat4-deficient mice. Nature 382, 174–177 (1996).

30. H. Baghdassarian, S. A. Blackstone, O. S. Clay, R. Philips, B. Matthiasardottir, M. Nehrebecky, V. K. Hua, R. McVicar, Y. Liu, S. M. Tucker, D. Randazzo, N. Deuitch, S. Rosenzweig, A. Mark, R. Sasik, K. M. Fisch, P. P. Chavan, E. Eren, N. R. Watts, C. A. Ma, M. Gadina, D. M. Schwartz, A. Sanyal, G. Werner, D. R. Murdock, N. Horita, S. Chowdhury, D. Dimmock, K. Jepsen, E. F. Remmers, R. Goldbach-Mansky, W. A. Gahl, J. J. O’Shea, J. D. Milner, N. E. Lewis, J. Chang, D. L. Kastner, K. Torok, H. Oda, C. D. Putnam, L. Broderick, Variant STAT4 and Response to Ruxolitinib in an Autoinflammatory Syndrome. New England Journal of Medicine 388, 2241–2252 (2023).

31. E. Xing, F. Ma, R. Wasikowski, A. C. Billi, M. Gharaee-Kermani, J. Fox, C. Dobry, A. Victory, M. K. Sarkar, X. Xing, O. Plazyo, H. W. Chen, G. Barber, H. Jacobe, P.-S. Tsou, R. L. Modlin, J. Varga, J. M. Kahlenberg, L. C. Tsoi, J. E. Gudjonsson, D. Khanna, Pansclerotic morphea is characterized by IFN-γ responses priming dendritic cell fibroblast crosstalk to promote fibrosis. JCI Insight 8, e171307 (2023).

32. D. J. Stein, M. Tran, I. Korsunsky, Accurate tiling of spatial single-cell data with Tessera. Bioinformatics [Preprint] (2025). 10.1101/2025.01.17.633630.

33. L. Steele, B. Olabi, K. Roberts, P. V. Mazin, S. Koplev, C. Tudor, B. Rumney, C. Admane, T. Jiang, D. Correa-Gallegos, K. P. Chakala, A. Binkevich, N. H. Gopee, A. Predeus, M. Prete, E. Winheim, K. Annusver, A. Forsthuber, L. Francis, S. Frech, C. Ganier, T. Layton, Y. Liu, H. Yuan, J. E. Gudjonsson, B. M. Lichtenberger, S. Mahil, J. Nanchahal, E. A. O’Toole, M. V. Plikus, Y. Rinkevich, E. Rognoni, C. H. Smith, S. A. Teichmann, M. Kasper, A. R. Foster, M. Lotfollahi, M. Haniffa, A single-cell and spatial genomics atlas of human skin fibroblasts reveals shared disease-related fibroblast subtypes across tissues. Nat Immunol 26, 1807–1820 (2025).

34. G. Werner, A. Sanyal, E. Mirizio, T. Hutchins, T. Tabib, R. Lafyatis, H. Jacobe, K. S. Torok, Single-Cell Transcriptome Analysis Identifies Subclusters with Inflammatory Fibroblast Responses in Localized Scleroderma. Int J Mol Sci 24, 9796 (2023).

35. Y. Gao, J. Li, W. Cheng, T. Diao, H. Liu, Y. Bo, C. Liu, W. Zhou, M. Chen, Y. Zhang, Z. Liu, W. Han, R. Chen, J. Peng, L. Zhu, W. Hou, Z. Zhang, Cross-tissue human fibroblast atlas reveals myofibroblast subtypes with distinct roles in immune modulation. Cancer Cell 42, 1764–1783.e10 (2024).

36. J. Sowerby, J. Choi, D. A. Rao, T Peripheral Helper Cells in Lymphoid Aggregate and Tertiary Lymphoid Structure Formation. Immunological Reviews 337, e70088 (2026).

37. D. Masopust, A. Awasthi, R. Bosselut, D. G. Brooks, M. Buggert, K. Chamoto, W. Cui, C. Dong, D. L. Farber, T. Gebhardt, C. Gerlach, A. Goldrath, P. D. Greenberg, J. S. Hale, A. Hayday, D. Homann, M. Iannacone, S. C. Jameson, M. K. Jenkins, N. S. Joshi, S. M. Kaech, A. Kallies, A. O. Kamphorst, M. H. Kaplan, P. Klenerman, M. Künzli, A. Lanzavecchia, G. M. Lauer, E. Lugli, A. D. Luster, L. K. Mackay, M. J. McElrath, S. N. Mueller, Z. Ndhlovu, T. Ndung’u, P. S. Ohashi, A. Oxenius, G. Pantaleo, M. Pepper, L. J. Picker, C. F. Quarnstrom, G. Reyes-Terán, M. Roederer, P. C. Rosato, G. S.-M. de Oca, F. Sallusto, T. N. Schumacher, D. M. Schwartz, E.-C. Shin, A. G. Soerens, D. S. Thommen, V. Vezys, J. P. B. Viola, B. D. Walker, T. H. Watts, C. T. Weaver, E. J. Wherry, H.-H. Xue, B. Youngblood, R. Ahmed, Guidelines for T cell nomenclature. Nat Rev Immunol 26, 298–313 (2026).

38. D. A. Mogilenko, O. Shpynov, P. S. Andhey, L. Arthur, A. Swain, E. Esaulova, S. Brioschi, I. Shchukina, M. Kerndl, M. Bambouskova, Z. Yao, A. Laha, K. Zaitsev, S. Burdess, S. Gillfilan, S. A. Stewart, M. Colonna, M. N. Artyomov, Comprehensive Profiling of an Aging Immune System Reveals Clonal GZMK+ CD8+ T Cells as Conserved Hallmark of Inflammaging. Immunity 54, 99–115.e12 (2021).

39. A. H. Jonsson, F. Zhang, G. Dunlap, E. Gomez-Rivas, G. F. M. Watts, H. J. Faust, K. V. Rupani, J. R. Mears, N. Meednu, R. Wang, G. Keras, J. S. Coblyn, E. M. Massarotti, D. J. Todd, J. H. Anolik, A. McDavid, Accelerating Medicines Partnership RA/SLE Network, K. Wei, D. A. Rao, S. Raychaudhuri, M. B. Brenner, Granzyme K ^+^ CD8 T cells form a core population in inflamed human tissue. Sci. Transl. Med. 14, eabo0686 (2022).

40. F. Lan, J. Li, W. Miao, F. Sun, S. Duan, Y. Song, J. Yao, X. Wang, C. Wang, X. Liu, J. Wang, L. Zhang, H. Qi, GZMK-expressing CD8+ T cells promote recurrent airway inflammatory diseases. Nature 638, 490–498 (2025).

41. I. C. Boothby, M. J. Kinet, D. P. Boda, E. Y. Kwan, S. Clancy, J. N. Cohen, I. Habrylo, M. M. Lowe, M. Pauli, A. E. Yates, J. D. Chan, H. W. Harris, I. M. Neuhaus, T. H. McCalmont, A. B. Molofsky, M. D. Rosenblum, Early-life inflammation primes a T helper 2 cell-fibroblast niche in skin. Nature 599, 667–672 (2021).

42. C. Gur, S.-Y. Wang, F. Sheban, M. Zada, B. Li, F. Kharouf, H. Peleg, S. Aamar, A. Yalin, D. Kirschenbaum, Y. Braun-Moscovici, D. A. Jaitin, T. Meir-Salame, E. Hagai, B. K. Kragesteen, B. Avni, S. Grisariu, C. Bornstein, S. Shlomi-Loubaton, E. David, R. Shreberk-Hassidim, V. Molho-Pessach, D. Amar, T. Tzur, R. Kuint, M. Gross, O. Barboy, A. Moshe, L. Fellus-Alyagor, D. Hirsch, Y. Addadi, S. Erenfeld, M. Biton, T. Tzemach, A. Elazary, Y. Naparstek, R. Tzemach, A. Weiner, A. Giladi, A. Balbir-Gurman, I. Amit, LGR5 expressing skin fibroblasts define a major cellular hub perturbed in scleroderma. Cell 185, 1373–1388.e20 (2022).

43. P. Fuschiotti, T. A. Medsger Jr., P. A. Morel, Effector CD8+ T cells in systemic sclerosis patients produce abnormally high levels of interleukin-13 associated with increased skin fibrosis. Arthritis & Rheumatism 60, 1119–1128 (2009).

44. P. Fuschiotti, A. T. Larregina, J. Ho, C. Feghali-Bostwick, T. A. Medsger Jr., Interleukin-13–producing CD8+ T cells mediate dermal fibrosis in patients with systemic sclerosis. Arthritis & Rheumatism 65, 236–246 (2013).

45. G. Li, A. T. Larregina, R. T. Domsic, D. B. Stolz, T. A. Medsger, R. Lafyatis, P. Fuschiotti, Skin-Resident Effector Memory CD8+CD28– T Cells Exhibit a Profibrotic Phenotype in Patients with Systemic Sclerosis. Journal of Investigative Dermatology 137, 1042–1050 (2017).

46. A. M. Gaydosik, T. Tabib, J. Das, A. Larregina, R. Lafyatis, P. Fuschiotti, Dysfunctional KLRB1+CD8+ T-cell responses are generated in chronically inflamed systemic sclerosis skin. Annals of the Rheumatic Diseases 84, 798–809 (2025).

47. E. Caves, A. Jussila, M. F. Forni, A. Benvie, V. Lei, D. King, H. Edelman, M. Hamdan, I. D. Odell, M. Hinchcliff, R. Atit, V. Horsley, *Atgl*-Dependent Adipocyte Lipolysis Promotes Lipodystrophy and Restrains Fibrogenic Responses during Skin Fibrosis. Journal of Investigative Dermatology 145, 1896–1909.e5 (2025).

48. L. Abbas, A. Joseph, E. Kunzler, H. T. Jacobe, Morphea: progress to date and the road ahead. Annals of Translational Medicine 9, 437–437 (2021).

49. Y. Rinkevich, G. G. Walmsley, M. S. Hu, Z. N. Maan, A. M. Newman, M. Drukker, M. Januszyk, G. W. Krampitz, G. C. Gurtner, H. P. Lorenz, I. L. Weissman, M. T. Longaker, Identification and isolation of a dermal lineage with intrinsic fibrogenic potential. Science 348, aaa2151 (2015).

50. M. B. Buechler, R. N. Pradhan, A. T. Krishnamurty, C. Cox, A. K. Calviello, A. W. Wang, Y. A. Yang, L. Tam, R. Caothien, M. Roose-Girma, Z. Modrusan, J. R. Arron, R. Bourgon, S. Müller, S. J. Turley, Cross-tissue organization of the fibroblast lineage. Nature 593, 575–579 (2021).

51. T. Tsukui, P. J. Wolters, D. Sheppard, Alveolar fibroblast lineage orchestrates lung inflammation and fibrosis. Nature 631, 627–634 (2024).

52. B. Hinz, D. Lagares, Evasion of apoptosis by myofibroblasts: a hallmark of fibrotic diseases. Nat Rev Rheumatol 16, 11–31 (2020).

53. N. G. Frangogiannis, Transforming growth factor–β in tissue fibrosis. J Exp Med 217, e20190103 (2020).

54. A. Prasse, D. V. Pechkovsky, G. B. Toews, W. Jungraithmayr, F. Kollert, T. Goldmann, E. Vollmer, J. Müller-Quernheim, G. Zissel, A vicious circle of alveolar macrophages and fibroblasts perpetuates pulmonary fibrosis via CCL18. Am J Respir Crit Care Med 173, 781–792 (2006).

55. P. Laurent, J. Lapoirie, D. Leleu, E. Levionnois, C. Grenier, B. Jurado-Mestre, E. Lazaro, P. Duffau, C. Richez, J. Seneschal, J.-L. Pellegrin, J. Constans, T. Schaeverbeke, I. Douchet, D. Duluc, T. Pradeu, C. Chizzolini, P. Blanco, M.-E. Truchetet, C. Contin-Bordes, Fédération Hospitalo-Universitaire ACRONIM and the Centre National de Référence des Maladies Auto-Immunes Systémiques Rares de l’Est et du Sud-Ouest (RESO), Interleukin-1β-Activated Microvascular Endothelial Cells Promote DC-SIGN-Positive Alternatively Activated Macrophages as a Mechanism of Skin Fibrosis in Systemic Sclerosis. Arthritis Rheumatol 74, 1013–1026 (2022).

56. C. A. Donado, E. Theisen, F. Zhang, A. Nathan, M. L. Fairfield, K. V. Rupani, D. Jones, K. P. Johannes, S. Raychaudhuri, D. F. Dwyer, A. H. Jonsson, M. B. Brenner, Granzyme K activates the entire complement cascade. Nature 641, 211–221 (2025).

57. D. Khanna, C. Padilla, L. C. Tsoi, V. Nagaraja, P. P. Khanna, T. Tabib, J. M. Kahlenberg, A. Young, S. Huang, J. E. Gudjonsson, D. A. Fox, R. Lafyatis, Tofacitinib blocks IFN-regulated biomarker genes in skin fibroblasts and keratinocytes in a systemic sclerosis trial. JCI Insight 7, e159566 (2022).

58. F. Chen, W. Ye, Q. Wang, L. Zhao, M. Liang, S. Zheng, T. Zhao, D. Xuan, Z. Zhu, Y. Yu, N. Kong, L. Jiang, X. Yang, X. Zhu, W. Wan, H. Zou, Y. Xue, BAricitinib in patients with SystemIC Sclerosis (BASICS): a prospective, open-label, randomised trial. Clin Rheumatol 44, 2861–2871 (2025).

59. A. Allawzi, H. Elajaili, E. F. Redente, E. Nozik-Grayck, Oxidative toxicology of bleomycin: Role of the extracellular redox environment. Current Opinion in Toxicology 13, 68–73 (2019).

60. L. A. Kalekar, J. N. Cohen, N. Prevel, P. M. Sandoval, A. N. Mathur, J. M. Moreau, M. M. Lowe, A. Nosbaum, P. J. Wolters, A. Haemel, F. Boin, M. D. Rosenblum, Regulatory T cells in skin are uniquely poised to suppress profibrotic immune responses. Sci Immunol 4, eaaw2910 (2019).

61. K. Bhamidipati, A. B. R. McIntyre, S. Kazerounian, G. Ce, S. W. Wong, M. Tran, S. A. Prell, R. Lau, V. Khedgikar, C. Altmann, A. Small, R. Madhu, S. R. Presti, K. S. Anufrieva, P. E. Blazar, J. K. Lange, J. A. Seifert, L. W. Moreland, A. P. Croft, M. H. Smith, L. T. Donlin, M. J. Lewis, A. H. Jonsson, C. Pitzalis, R. Thomas, E. M. Gravallese, M. B. Brenner, I. Korsunsky, M. D. Wechalekar, K. Wei, Spatial patterning of fibroblast TGFβ signaling underlies treatment resistance in rheumatoid arthritis. Nat Immunol 27, 556–571 (2026).

62. D. M. Cable, E. Murray, L. S. Zou, A. Goeva, E. Z. Macosko, F. Chen, R. A. Irizarry, Robust decomposition of cell type mixtures in spatial transcriptomics. Nat Biotechnol 40, 517–526 (2022).

63. M. R. Vahid, E. L. Brown, C. B. Steen, W. Zhang, H. S. Jeon, M. Kang, A. J. Gentles, A. M. Newman, High-resolution alignment of single-cell and spatial transcriptomes with CytoSPACE. Nat Biotechnol 41, 1543–1548 (2023).

64. S. Jin, M. V. Plikus, Q. Nie, CellChat for systematic analysis of cell–cell communication from single-cell transcriptomics. Nat Protoc 20, 180–219 (2025).

65. J. Sbierski-Kind, K. M. Cautivo, J. Nilsson, J. C. Wagner, M. W. Dahlgren, N. E. Crystal, M. McClave, N. M. Mroz, M. Ganslmeier, C. O. Lizama, A. L. Gan, P. R. Matatia, M. T. Taruselli, A. A. Chang, S. Caryotakis, C. E. O’Leary, M. Kotas, J.-H. Lee, T. Gu, H. Seo, H. J. Kim, A. N. Mattis, T. Peng, R. M. Locksley, A. B. Molofsky, Type 2 lymphocytes restrict type 3 lymphocytes during liver fibrosis and colocalize in fibroblast niches. Science Advances 12, eaea6805 (2026).

