## Supplementary materials 1 for "Spatial Orchestration of Skin Fibrosis by a CD8+ T cell-Myofibroblast Axis"

**by Boothby IC, *et al.***

Figures S1-S8

Tables S1-S2

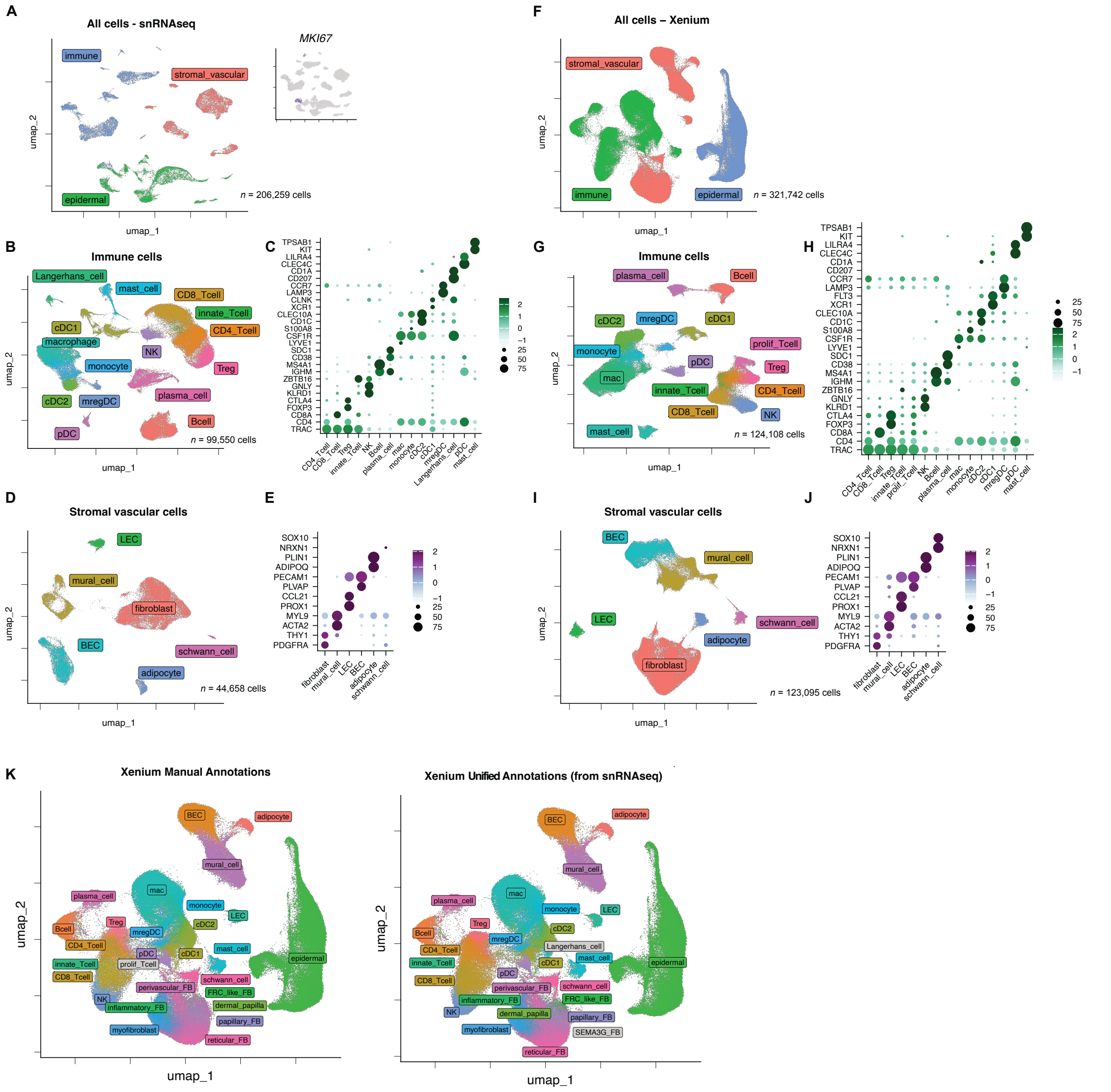


Fig. S1. **A multimodal atlas of morphea and eosinophilic fasciitis skin. (A-E)** UMAP plots and expression of selected marker genes in immune and stromal populations in snRNAseq data from healthy, morphea, and eosinophilic fasciitis samples. **(F-I)** UMAP plots and expression of selected marker genes in immune and stromal populations in Xenium data. **(K)** Comparison of manual annotations of Xenium data to multimodal annotations generated using diffusion-based label propagation from snRNAseq reference annotations.

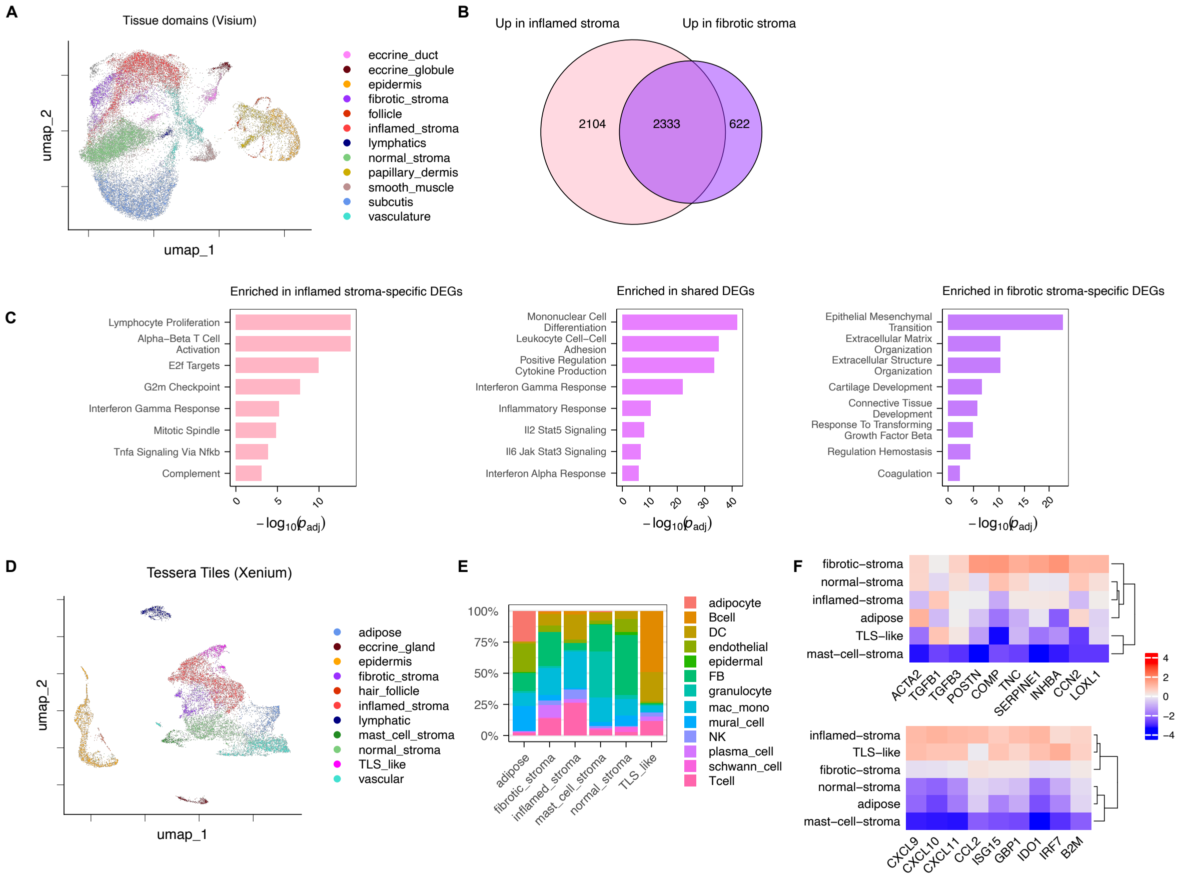


**Fig. S2. Fibrosing skin is zonated into inflammatory and fibrotic domains. (A)** Full UMAP plot of Visium tissue domains. **(B)** Pseudobulk differential gene expression testing was performed between inflamed vs. normal stromal domains and fibrotic vs. normal domains. Overlap of differentially expressed genes (DEGs) is shown. **(C)** Enrichment of gene ontology and MsigDB Hallmark terms within inflamed-exclusive, fibrotic-exclusive, and shared DEG lists. **(D)** UMAP of tissue domains generated from Xenium data using Tessera. **(E)** Frequency of immune and stromal cell types in Xenium tissue domains. **(F)** Expression of fibrosis-associated and IFN-γ stimulated genes in Xenium tissue domains.

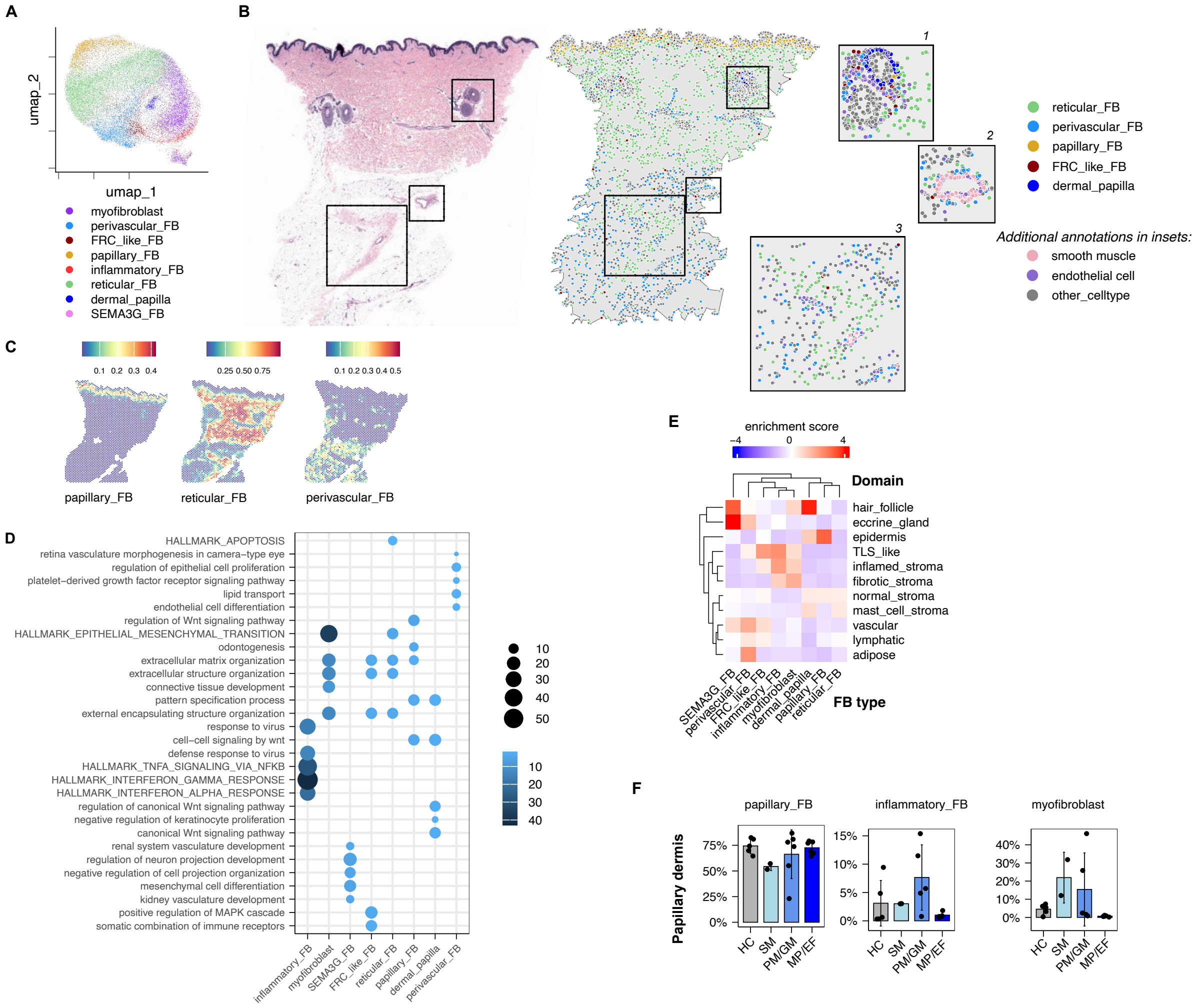


**Fig. S3. Localization and function of fibroblast subsets in healthy and fibrotic skin.**

**(A)** UMAP of fibroblast from Xenium data, annotated using labels transferred from snRNAseq. **(B)** H&E and Xenium images of a healthy skin sample demonstrating localization of select fibroblast subsets. *Inset 1:* perifollicular fibroblasts. *Inset 2:* perivascular fibroblasts. *Inset 3:* fibrous septa of the subcutis. **(C)** Predicted fibroblast cell type frequencies in the same skin sample, analyzed using Visium. **(D)** Enrichment of MsigDB hallmark and gene ontology signatures in each fibroblast subset. **(E)** Enrichment of fibroblast subtypes in Xenium spatial domains. Higher enrichment scores indicate a cell type is more likely to be found in a spatial domain than expected by chance. **(F)** Per-individual quantification of fibroblast subsets in manually annotated papillary dermis (accompanying fig. 3E).

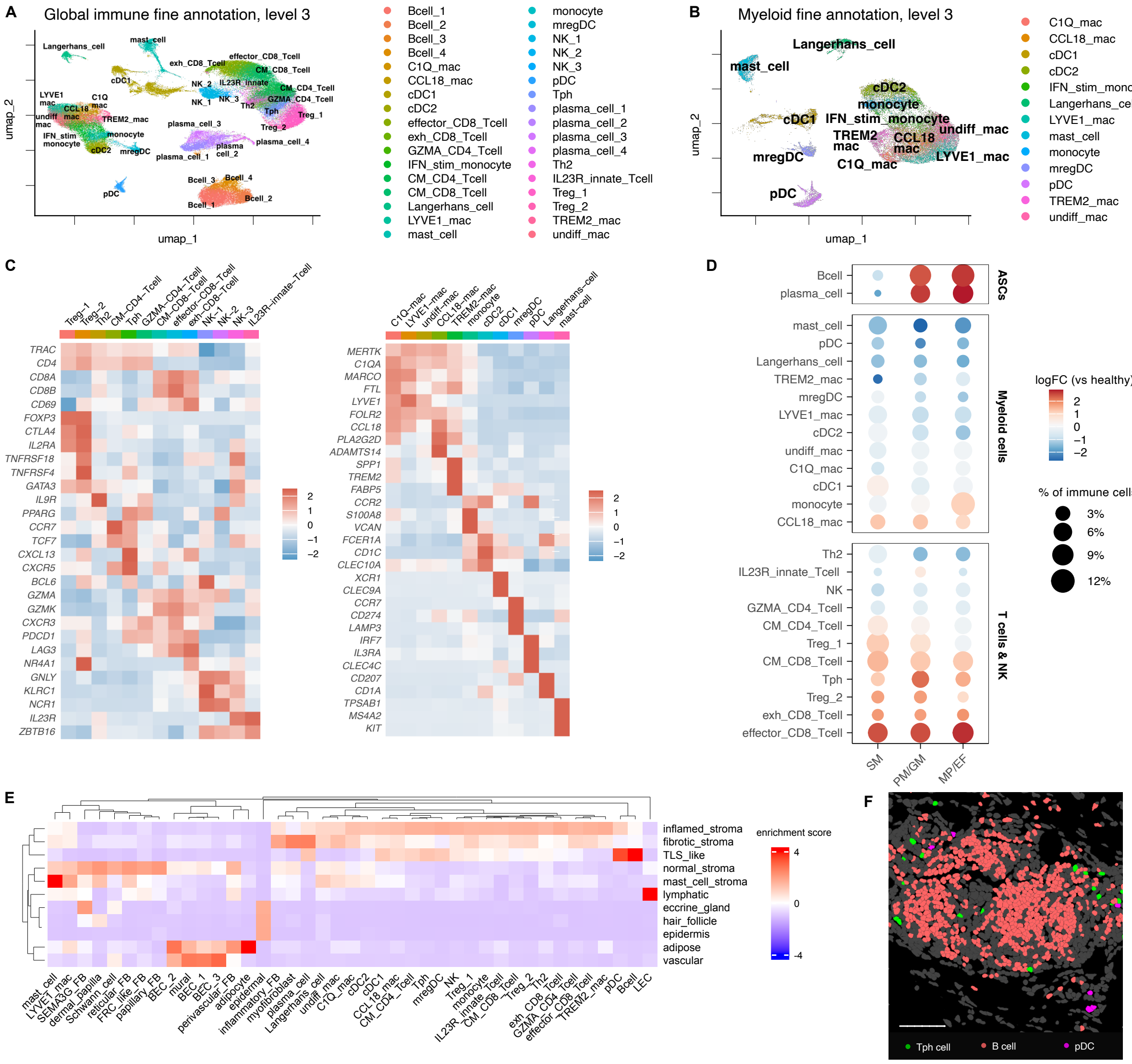


**Fig. S4. Immune composition of fibrotic skin. (A)** UMAP of level 3 fine annotation of all immune cell subsets from snRNAseq data. **(B)** UMAP of level 3 fine annotation of myeloid subsets from snRNAseq data. **(C)** Heatmap of marker gene expression in T cell subsets (left) and myeloid cell subsets (right) in snRNAseq data. **(D)** Immune cell abundance in each disease subtype. Color scale represents the change in immune cell abundance relative to healthy skin. **(E)** Enrichment of immune cells in Xenium spatial domains. Higher enrichment scores indicate a cell type is more likely to be found in a spatial domain than expected by chance. **(F)** Representative Xenium images of tertiary lymphoid structure-like immune cell clusters in plaque morphea skin. *Scale bar*: 100µm.

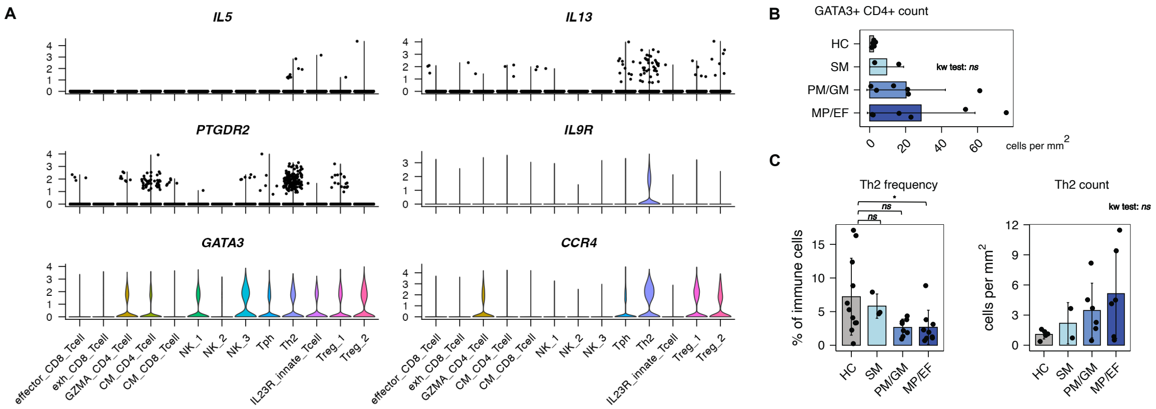


**Fig. S5. Th2 cells are not preferentially increased in morphea and EF. (A)** Violin plots of expression of Th2-associated genes in snRNAseq data. **(B)** Count of all GATA3+ CD4+ cells in Xenium data. **(C)** Frequency of Th2 cells in snRNAseq data and Th2 counts in Xenium data. Statistics calculated with Kruskall-Wallis test with post-hoc Dunn testing. *All plots:* **p* < 0.05; ***p* < 0.05; ****p* < 0.005; *ns*, not significant.

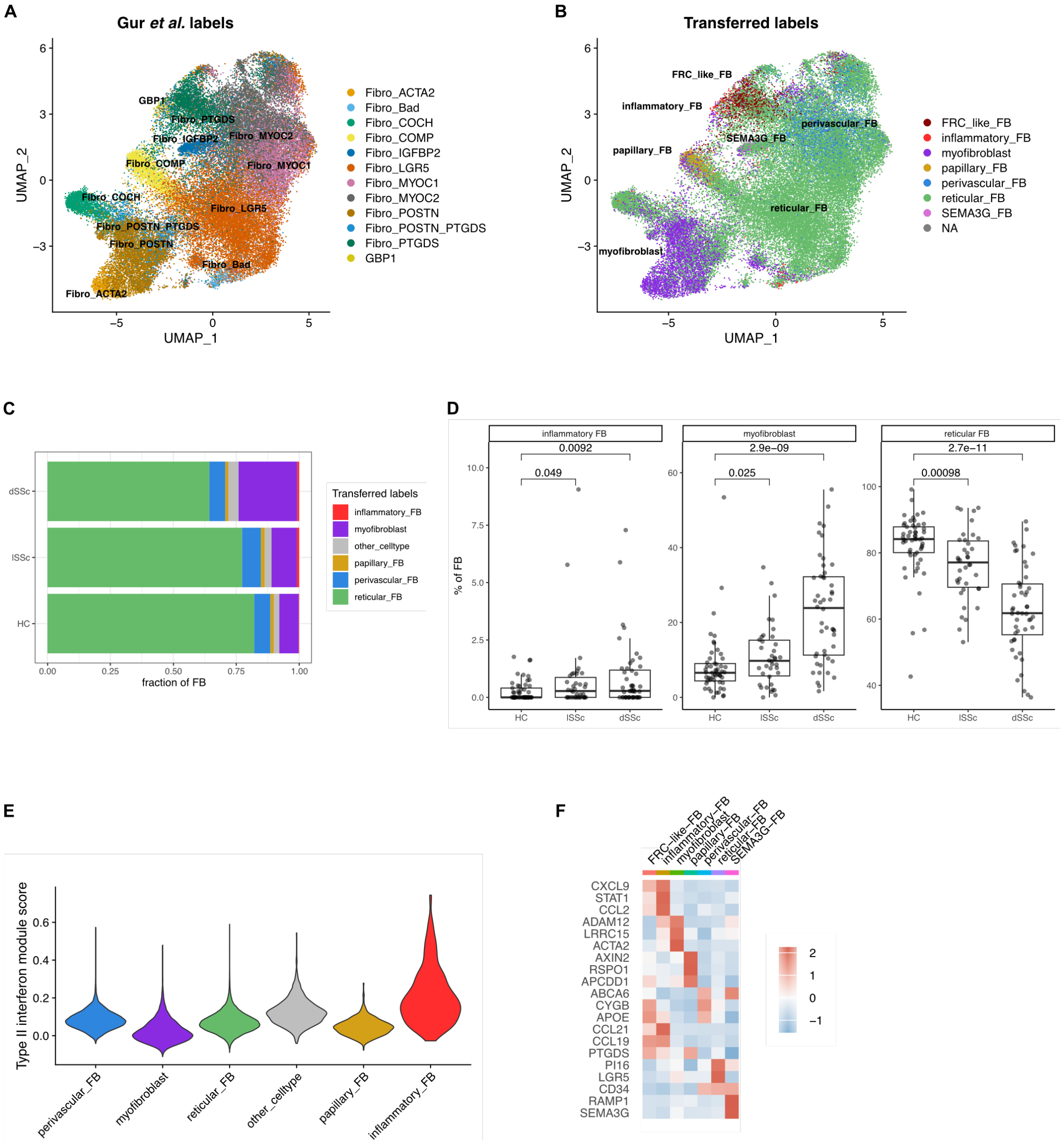


**Figure S6- SSc skin exhibits replacement of reticular fibroblasts by inflammatory fibroblasts and myofibroblasts. (A)** UMAP of fibroblasts from SSc and healthy control skin with annotations from original publication. **(B)** UMAP of fibroblasts annotated using labels transferred from fibroblast subsets described in Fig 2. **(C)** Proportion of indicated fibroblast subset among total fibroblasts for all patients in each clinical category. **(D)** Proportion of indicated fibroblast subsets among total fibroblasts, per patient, grouped by clinical category. Boxes indicate median and interquartile range (IQR); whiskers extend to the most extreme data point within 1.5 IQR of the box edges. Statistical comparisons by Wilcoxon rank-sum test with exact p-values shown. **(E)** Type II interferon module score for each fibroblast subset using Seurat AddModuleScore function. **(F)** Dotplot showing expression of genes making up type II ISG in each fibroblast subset.

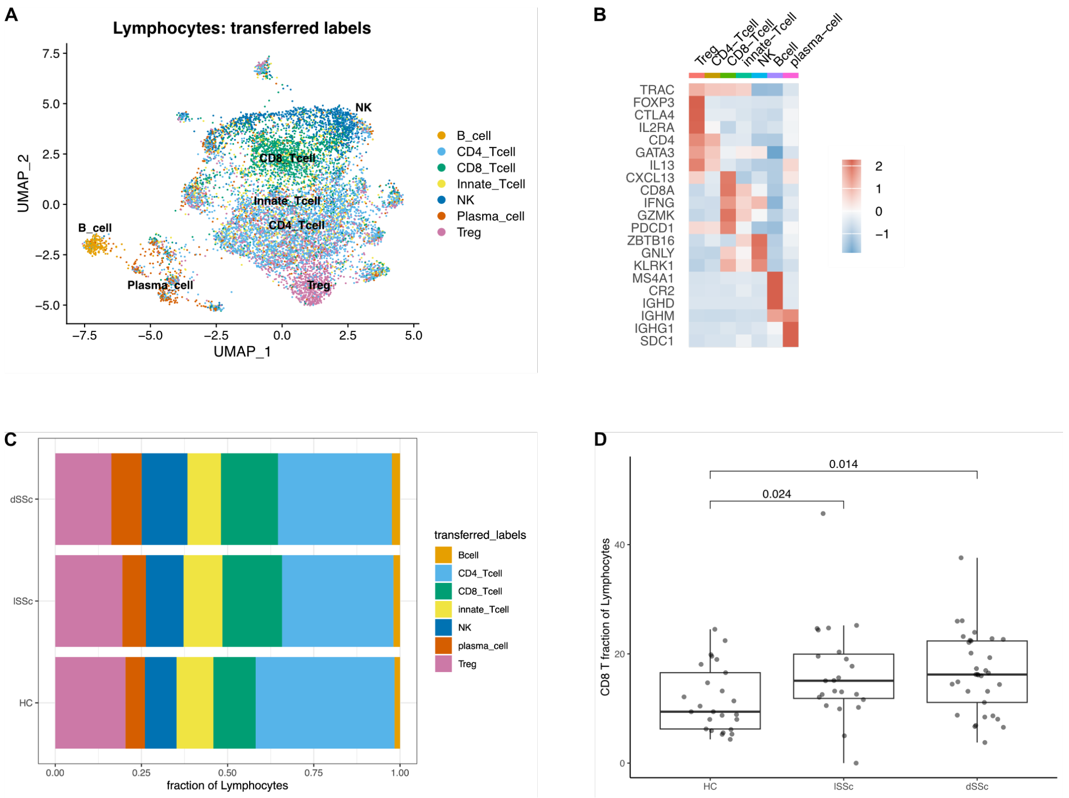


**Fig. S7. SSc skin exhibits expanded CD8+ T cells. (A)** UMAP of skin immune cells from SSc and healthy control skin with annotations from original publication. **(B)** UMAP of skin immune cells annotated using labels transferred from cell types described in. **(C)** Feature plot of *CD8A* expression in skin immune UMAP. **(D)** Proportion of CD8+ T cells among total immune cells, per patient, grouped by clinical category. **(E)** Proportion of CD8+ T cells among total immune cells, per patient, grouped by Scl70 antibody status. For (D) and (E), boxes indicate median and interquartile range (IQR); whiskers extend to the most extreme data point within 1.5 IQR of the box edges; statistical comparisons by Wilcoxon rank-sum test with exact p-values shown.

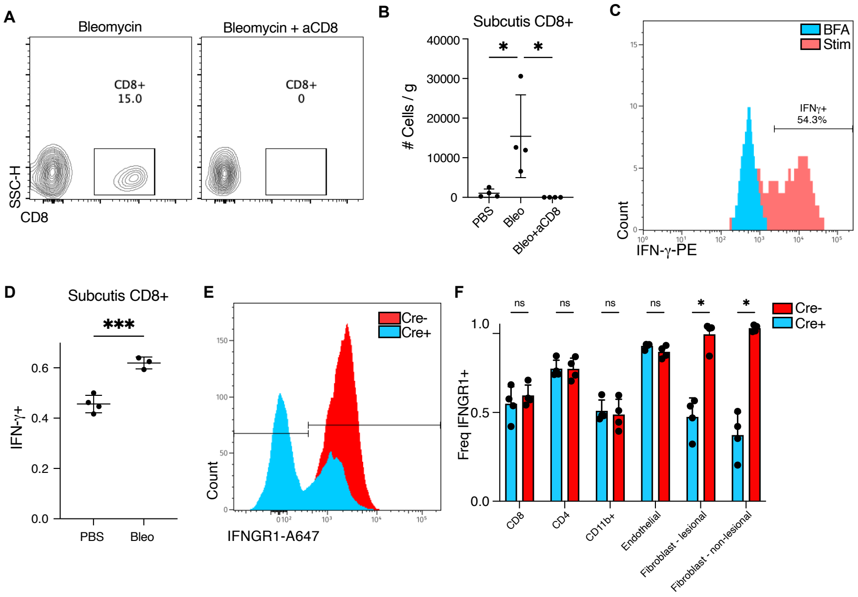


**Fig. S8. Manipulation of CD8+ T cells and fibroblast IFNγ-R in mouse skin fibrosis.**

**(A)** Representative skin flow cytometry plots of skin CD8+ T cells isolated from mice treated with bleomycin vs bleomycin + α-CD8 after 7 days, pregated on live/CD45+. **(B)** Flow cytometric enumeration of CD8+ T cells in mechanically separated subcutis layers of bleomycin or control (PBS) skin lesions, normalized to skin weight. Statistics calculated using one-way ANOVA with Šidák's multiple comparisons adjustment. **(C)** Representative histogram of IFN-γ staining in *in vitro* re-stimulated subcutis CD8+ T cells, pre-gated as live/CD45+/CD11b-/Thy1+/CD3+/TCRβ+ cells, with unstimulated (BFA) control shown. **(D)** Proportion of IFN-γ+ cells among subcutis CD8+ T cells from indicated skin treatments. Statistics calculated using unpaired Welch's t-test. **(E-F)** Flow cytometry of $Ifngr1^{fl/fl}$ mice with and without $Pdgfra^{CreER}$. All mice were treated with tamoxifen. Representative histograms of IFNGR1 staining in live/CD45-/CD31-/PDPN+/PDGFRα+ skin fibroblasts (E), quantification of IFNGR1+ cells in immune and stromal subsets (F). Data in B, D, F are representative of two independent experiments with similar results. Statistics calculated using unpaired t-tests with Welch's correction and Holm-Šidák's multiple comparisons adjustment. In all plots, *p < 0.05; **p < 0.01; ***p < 0.005; ns, not significant.

**Table S1. Roster of patient samples.**

| Patient ID | Diagnosis | Supergroup | Pathology Stage | Age | Sex | Skin Site |
|---|---|---|---|---|---|---|
| HS4 | Healthy control | HC | healthy | 72 | F | thigh |
| HS5 | Healthy control | HC | healthy | 62 | M | leg |
| HS6 | Healthy control | HC | healthy | 64 | M | leg |
| HS24 | Healthy control | HC | healthy | 64 | M | arm |
| HS30 | Healthy control | HC | healthy | 63 | M | abdomen |
| HS31 | Healthy control | HC | healthy | 41 | F | leg |
| HS32 | Healthy control | HC | healthy | 44 | F | arm |
| HS33 | Healthy control | HC | healthy | 71 | F | leg |
| HS34 | Healthy control | HC | healthy | 85 | F | leg |
| HS35 | Healthy control | HC | healthy | 67 | M | arm |
| HS36 | Healthy control | HC | healthy | 55 | F | arm |
| HS10 | Superficial morphea | SM | inflammatory | 45 | F | abdomen |
| HS14 | Superficial morphea | SM | inflammatory | 66 | F | abdomen |
| HS20 | Superficial morphea | SM | early | 69 | F | shoulder |
| HS11 | Plaque morphea | PM/GM | inflammatory | 75 | F | breast |
| HS16 | Generalized morphea | PM/GM | inflammatory | 66 | F | breast |
| HS2 | Generalized morphea | PM/GM | late | 88 | F | posterior thigh |
| HS13 | Generalized morphea | PM/GM | inflammatory | 74 | F | breast |
| HS18 | Plaque morphea | PM/GM | inflammatory | 59 | M | forearm |
| HS15 | Plaque morphea | PM/GM | late | 58 | F | abdomen |
| HS19 | Plaque morphea | PM/GM | inflammatory | 87 | F | breast |
| HS22 | Generalized morphea | PM/GM | inflammatory | 69 | F | abdomen |
| HS9 | Eosinophilic fasciitis | MP/EF | inflammatory | 44 | F | upper arm |
| HS8 | Eosinophilic fasciitis | MP/EF | inflammatory | 58 | M | upper arm |
| HS1 | Eosinophilic fasciitis | MP/EF | late | 74 | F | anterior lower leg |
| HS17 | Morphea profunda | MP/EF | inflammatory | 19 | F | abdomen |
| HS12 | Morphea profunda | MP/EF | inflammatory | 44 | F | upper arm |
| HS25 | Eosinophilic fasciitis | MP/EF | inflammatory | 61 | F | forearm |
| HS26 | Eosinophilic fasciitis | MP/EF | late | 71 | M | leg |
| HS27 | Morphea profunda | MP/EF | late | 47 | M | abdomen |
| HS28 | Eosinophilic fasciitis | MP/EF | inflammatory | 52 | F | forearm |
| HS21 | Linear morphea | NA | inflammatory | 37 | M | upper arm |
