## Supplementary material for "Spatial Orchestration of Skin Fibrosis by a CD8+ T cell-Myofibroblast Axis": Table S2

**Table S2. List of genes in custom Xenium probeset**

| Gene | Ensembl ID | Probesets |
|---|---|---|
| C3 | ENSG00000125730 | 3 |
| KRT14 | ENSG00000186847 | 3 |
| HLA-DPA1 | ENSG00000231389 | 4 |
| HLA-DQA1 | ENSG00000196735 | 4 |
| HLA-DRA | ENSG00000204287 | 4 |
| IL9 | ENSG00000145839 | 6 |
| ACSBG1 | ENSG00000103740 | 7 |
| ACTA2 | ENSG00000107796 | 3 |
| ADIPOQ | ENSG00000181092 | 5 |
| APOD | ENSG00000189058 | 3 |
| B2M | ENSG00000166710 | 6 |
| C1QA | ENSG00000173372 | 7 |
| C1QB | ENSG00000173369 | 5 |
| C1QC | ENSG00000159189 | 7 |
| CCN2 | ENSG00000118523 | 4 |
| CD163 | ENSG00000177575 | 3 |
| CFD | ENSG00000197766 | 4 |
| CFH | ENSG00000000971 | 3 |
| COMP | ENSG00000105664 | 4 |
| CTHRC1 | ENSG00000164932 | 6 |
| CXCL14 | ENSG00000145824 | 4 |
| FABP4 | ENSG00000170323 | 5 |
| HDC | ENSG00000140287 | 7 |
| IGFBP7 | ENSG00000163453 | 4 |
| IGHM | ENSG00000211899 | 7 |
| KRT15 | ENSG00000171346 | 6 |
| KRTDAP | ENSG00000188508 | 3 |
| LCE2B | ENSG00000159455 | 6 |
| LYZ | ENSG00000090382 | 4 |
| MCAM | ENSG00000076706 | 5 |
| MYH11 | ENSG00000133392 | 6 |
| MYL9 | ENSG00000101335 | 5 |
| NOTCH3 | ENSG00000074181 | 7 |
| PI16 | ENSG00000164530 | 5 |
| PLIN1 | ENSG00000166819 | 4 |
| PNPLA2 | ENSG00000177666 | 5 |

| | | |
|---|---|---|
| POSTN | ENSG00000133110 | 3 |
| PTGDS | ENSG00000107317 | 6 |
| SERPINE1 | ENSG00000106366 | 5 |
| TIMP2 | ENSG00000035862 | 6 |
| TNC | ENSG00000041982 | 6 |
| TPSAB1 | ENSG00000172236 | 3 |
| TYRP1 | ENSG00000107165 | 6 |
| CD4 | ENSG00000010610 | 8 |
| GATA3 | ENSG00000107485 | 8 |
| RORA | ENSG00000069667 | 8 |
| RORC | ENSG00000143365 | 8 |
| TBX21 | ENSG00000073861 | 8 |
| PPARG | ENSG00000132170 | 8 |
| ACKR4 | ENSG00000129048 | 8 |
| CCN5 | ENSG00000064205 | 8 |
| DPP4 | ENSG00000197635 | 8 |
| GAS1 | ENSG00000180447 | 8 |
| ISLR | ENSG00000129009 | 8 |
| OGN | ENSG00000106809 | 8 |
| PTGIS | ENSG00000124212 | 8 |
| SFRP2 | ENSG00000145423 | 8 |
| SLPI | ENSG00000124107 | 7 |
| CD226 | ENSG00000150637 | 8 |
| CD28 | ENSG00000178562 | 8 |
| CD3E | ENSG00000198851 | 8 |
| CD69 | ENSG00000110848 | 8 |
| LCK | ENSG00000182866 | 8 |
| TRAC | ENSG00000277734 | 8 |
| TRBC1 | ENSG00000211751 | 7 |
| TRBC2 | ENSG00000211772 | 5 |
| TRDC | ENSG00000211829 | 8 |
| TRGC1 | ENSG00000211689 | 8 |
| TRGC2 | ENSG00000227191 | 8 |
| ZAP70 | ENSG00000115085 | 8 |
| BATF | ENSG00000156127 | 5 |
| CTLA4 | ENSG00000163599 | 8 |
| FOXP3 | ENSG00000049768 | 8 |
| JAG1 | ENSG00000101384 | 8 |
| LAYN | ENSG00000204381 | 8 |

| | | |
|---|---|---|
| LRRC32 | ENSG00000137507 | 8 |
| TNFRSF4 | ENSG00000186827 | 1 |
| TIGIT | ENSG00000181847 | 8 |
| CD27 | ENSG00000139193 | 8 |
| CDH11 | ENSG00000140937 | 8 |
| FAP | ENSG00000078098 | 8 |
| LAMA2 | ENSG00000196569 | 8 |
| LOXL1 | ENSG00000129038 | 8 |
| LTBP2 | ENSG00000119681 | 8 |
| MFAP5 | ENSG00000197614 | 8 |
| SFRP4 | ENSG00000106483 | 8 |
| SULF1 | ENSG00000137573 | 8 |
| TMEM119 | ENSG00000183160 | 8 |
| TNFRSF12A | ENSG00000006327 | 7 |
| ADAM12 | ENSG00000148848 | 8 |
| C7 | ENSG00000112936 | 8 |
| MYOC | ENSG00000034971 | 8 |
| NRP1 | ENSG00000099250 | 8 |
| RBP5 | ENSG00000139194 | 8 |
| RIPOR3 | ENSG00000042062 | 8 |
| CCL19 | ENSG00000172724 | 7 |
| CD8A | ENSG00000153563 | 8 |
| CD8B | ENSG00000172116 | 8 |
| GNLY | ENSG00000115523 | 8 |
| GZMB | ENSG00000100453 | 8 |
| GZMA | ENSG00000145649 | 8 |
| GZMK | ENSG00000113088 | 8 |
| LAG3 | ENSG00000089692 | 8 |
| KLRC3 | ENSG00000205810 | 8 |
| XCL1 | ENSG00000143184 | 8 |
| XCL2 | ENSG00000143185 | 6 |
| ABCA10 | ENSG00000154263 | 8 |
| ADAM33 | ENSG00000149451 | 8 |
| ADAMTS10 | ENSG00000142303 | 8 |
| DNM1 | ENSG00000106976 | 8 |
| GLI3 | ENSG00000106571 | 8 |
| PAPLN | ENSG00000100767 | 8 |
| PLAGL1 | ENSG00000118495 | 8 |
| PLEKHH2 | ENSG00000152527 | 8 |

| | | |
|---|---|---|
| SNED1 | ENSG00000162804 | 8 |
| SYNE1 | ENSG00000131018 | 8 |
| TRIP10 | ENSG00000125733 | 8 |
| TWIST2 | ENSG00000233608 | 8 |
| TWIST1 | ENSG00000122691 | 8 |
| SNAI2 | ENSG00000019549 | 8 |
| MMP11 | ENSG00000099953 | 8 |
| KRT16 | ENSG00000186832 | 8 |
| GBP1 | ENSG00000117228 | 8 |
| ISG15 | ENSG00000187608 | 6 |
| ITGAX | ENSG00000140678 | 8 |
| AICDA | ENSG00000111732 | 8 |
| BCL2 | ENSG00000171791 | 8 |
| MYC | ENSG00000136997 | 8 |
| CD209 | ENSG00000090659 | 8 |
| CSF1R | ENSG00000182578 | 8 |
| CX3CR1 | ENSG00000168329 | 8 |
| FOLR2 | ENSG00000165457 | 8 |
| LILRB2 | ENSG00000131042 | 8 |
| MARCO | ENSG00000019169 | 8 |
| MERTK | ENSG00000153208 | 8 |
| MRC1 | ENSG00000260314 | 8 |
| SELENOP | ENSG00000250722 | 8 |
| SIGLEC1 | ENSG00000088827 | 8 |
| TLR4 | ENSG00000136869 | 8 |
| LYVE1 | ENSG00000133800 | 8 |
| LILRB5 | ENSG00000105609 | 8 |
| SPP1 | ENSG00000118785 | 8 |
| CLEC10A | ENSG00000132514 | 8 |
| CXCL6 | ENSG00000124875 | 8 |
| CSMD2 | ENSG00000121904 | 8 |
| MMP1 | ENSG00000196611 | 8 |
| MMP3 | ENSG00000149968 | 8 |
| PCDH7 | ENSG00000169851 | 8 |
| SCARA5 | ENSG00000168079 | 8 |
| TENM3 | ENSG00000218336 | 8 |
| WNT2 | ENSG00000105989 | 8 |
| WNT5A | ENSG00000114251 | 8 |
| BCL6 | ENSG00000113916 | 8 |

| | | |
|---|---|---|
| CD200 | ENSG00000091972 | 8 |
| CXCR5 | ENSG00000160683 | 8 |
| CXCR6 | ENSG00000172215 | 8 |
| KLRB1 | ENSG00000111796 | 8 |
| TOX2 | ENSG00000124191 | 8 |
| PDCD1 | ENSG00000188389 | 8 |
| APOC1 | ENSG00000130208 | 6 |
| CD36 | ENSG00000135218 | 8 |
| FABP5 | ENSG00000164687 | 8 |
| MMP9 | ENSG00000100985 | 8 |
| TREM2 | ENSG00000095970 | 8 |
| APOE | ENSG00000130203 | 4 |
| CCR3 | ENSG00000183625 | 8 |
| EPX | ENSG00000121053 | 8 |
| MBP | ENSG00000197971 | 8 |
| SIGLEC8 | ENSG00000105366 | 8 |
| ANGPTL1 | ENSG00000116194 | 8 |
| CD55 | ENSG00000196352 | 8 |
| COL6A5 | ENSG00000172752 | 8 |
| PDGFA | ENSG00000197461 | 8 |
| PDGFB | ENSG00000100311 | 8 |
| PDGFC | ENSG00000145431 | 8 |
| PDGFD | ENSG00000170962 | 8 |
| PDGFRA | ENSG00000134853 | 8 |
| PDGFRB | ENSG00000113721 | 8 |
| THY1 | ENSG00000154096 | 8 |
| PTPRC | ENSG00000081237 | 8 |
| CD40 | ENSG00000101017 | 8 |
| FCER2 | ENSG00000104921 | 8 |
| IGHD | ENSG00000211898 | 8 |
| TCL1A | ENSG00000100721 | 8 |
| CD177 | ENSG00000204936 | 8 |
| CEACAM8 | ENSG00000124469 | 8 |
| CXCR1 | ENSG00000163464 | 8 |
| CXCR2 | ENSG00000180871 | 8 |
| MPO | ENSG00000005381 | 8 |
| CLEC4C | ENSG00000198178 | 8 |
| IL3RA | ENSG00000185291 | 8 |
| IRF7 | ENSG00000185507 | 8 |

| | | |
|---|---|---|
| LILRA4 | ENSG00000239961 | 8 |
| PLAC8 | ENSG00000145287 | 8 |
| SMPD3 | ENSG00000103056 | 8 |
| VIPR2 | ENSG00000106018 | 8 |
| AIM2 | ENSG00000163568 | 8 |
| CD24 | ENSG00000272398 | 8 |
| CR1 | ENSG00000203710 | 8 |
| CR2 | ENSG00000117322 | 8 |
| TNFRSF13B | ENSG00000240505 | 7 |
| GPD1 | ENSG00000167588 | 8 |
| LIPE | ENSG00000079435 | 8 |
| SAA1 | ENSG00000173432 | 8 |
| CLEC9A | ENSG00000197992 | 8 |
| HAVCR2 | ENSG00000135077 | 8 |
| IRF8 | ENSG00000140968 | 8 |
| THBD | ENSG00000178726 | 8 |
| XCR1 | ENSG00000173578 | 8 |
| FLT3 | ENSG00000122025 | 8 |
| CCR7 | ENSG00000126353 | 8 |
| CD274 | ENSG00000120217 | 8 |
| IDO1 | ENSG00000131203 | 8 |
| LAMP3 | ENSG00000078081 | 8 |
| PDCD1LG2 | ENSG00000197646 | 8 |
| ZBTB46 | ENSG00000130584 | 8 |
| IFNG | ENSG00000111537 | 8 |
| IL4R | ENSG00000077238 | 8 |
| IL6R | ENSG00000160712 | 8 |
| OSMR | ENSG00000145623 | 8 |
| TGFBR1 | ENSG00000106799 | 8 |
| TGFBR2 | ENSG00000163513 | 8 |
| IL23R | ENSG00000162594 | 8 |
| IL7R | ENSG00000168685 | 8 |
| IL2RA | ENSG00000134460 | 8 |
| IL5RA | ENSG00000091181 | 8 |
| IL11 | ENSG00000095752 | 8 |
| IL4 | ENSG00000113520 | 7 |
| IFNB1 | ENSG00000171855 | 8 |
| CXCL10 | ENSG00000169245 | 8 |
| CXCL11 | ENSG00000169248 | 8 |

| | | |
|---|---|---|
| IL33 | ENSG00000137033 | 8 |
| IL23A | ENSG00000110944 | 8 |
| OSM | ENSG00000099985 | 8 |
| TGFB1 | ENSG00000105329 | 8 |
| TGFB2 | ENSG00000092969 | 8 |
| TGFB3 | ENSG00000119699 | 8 |
| TNF | ENSG00000232810 | 8 |
| CD38 | ENSG00000004468 | 8 |
| CD79A | ENSG00000105369 | 8 |
| IRF4 | ENSG00000137265 | 8 |
| PRDM16 | ENSG00000142611 | 8 |
| SDC1 | ENSG00000115884 | 8 |
| SPAG4 | ENSG00000061656 | 8 |
| XBP1 | ENSG00000100219 | 8 |
| ZNF215 | ENSG00000149054 | 8 |
| CDK1 | ENSG00000170312 | 8 |
| MKI67 | ENSG00000148773 | 8 |
| TOP2A | ENSG00000131747 | 8 |
| CDKN2A | ENSG00000147889 | 8 |
| CDKN1A | ENSG00000124762 | 8 |
| CSF3R | ENSG00000119535 | 8 |
| FGL2 | ENSG00000127951 | 8 |
| RTN1 | ENSG00000139970 | 8 |
| TGFBI | ENSG00000120708 | 8 |
| CD1C | ENSG00000158481 | 8 |
| CD1D | ENSG00000158473 | 8 |
| FCER1A | ENSG00000179639 | 8 |
| CD1A | ENSG00000158477 | 8 |
| CD207 | ENSG00000116031 | 8 |
| CDH20 | ENSG00000101542 | 8 |
| CFAP73 | ENSG00000186710 | 8 |
| ADAM19 | ENSG00000135074 | 8 |
| CCL21 | ENSG00000137077 | 8 |
| FLT4 | ENSG00000037280 | 8 |
| MMRN1 | ENSG00000138722 | 8 |
| PROX1 | ENSG00000117707 | 8 |
| STAB2 | ENSG00000136011 | 8 |
| TFF3 | ENSG00000160180 | 8 |
| CD19 | ENSG00000177455 | 8 |

| | | |
|---|---|---|
| CD22 | ENSG00000012124 | 8 |
| CD79B | ENSG00000007312 | 8 |
| FCRL2 | ENSG00000132704 | 8 |
| MS4A1 | ENSG00000156738 | 8 |
| PAX5 | ENSG00000196092 | 8 |
| ABCA8 | ENSG00000141338 | 8 |
| APCDD1 | ENSG00000154856 | 8 |
| AXIN2 | ENSG00000168646 | 8 |
| COL13A1 | ENSG00000197467 | 8 |
| COL18A1 | ENSG00000182871 | 8 |
| NKD2 | ENSG00000145506 | 8 |
| RSPO1 | ENSG00000169218 | 8 |
| TBX18 | ENSG00000112837 | 8 |
| IL13 | ENSG00000169194 | 8 |
| IL15 | ENSG00000164136 | 8 |
| IL17A | ENSG00000112115 | 8 |
| IL21 | ENSG00000138684 | 8 |
| IL10 | ENSG00000136634 | 8 |
| AREG | ENSG00000109321 | 8 |
| CSF1 | ENSG00000184371 | 8 |
| LHX6 | ENSG00000106852 | 8 |
| PECAM1 | ENSG00000261371 | 8 |
| PLVAP | ENSG00000130300 | 8 |
| RAMP3 | ENSG00000122679 | 8 |
| TACR1 | ENSG00000115353 | 8 |
| PODXL | ENSG00000128567 | 8 |
| HEY1 | ENSG00000164683 | 8 |
| IGF2 | ENSG00000167244 | 8 |
| JAG2 | ENSG00000184916 | 8 |
| MECOM | ENSG00000085276 | 8 |
| SEMA3G | ENSG00000010319 | 8 |
| SOX17 | ENSG00000164736 | 8 |
| FGFR1 | ENSG00000077782 | 8 |
| GREM1 | ENSG00000166923 | 8 |
| LGR6 | ENSG00000133067 | 8 |
| TBX1 | ENSG00000184058 | 8 |
| TRPM3 | ENSG00000083067 | 8 |
| WIF1 | ENSG00000156076 | 8 |
| LGR5 | ENSG00000139292 | 8 |

| | | |
|---|---|---|
| GATA6 | ENSG00000141448 | 8 |
| KRT6A | ENSG00000205420 | 8 |
| KRT6B | ENSG00000185479 | 8 |
| PTN | ENSG00000105894 | 8 |
| RBP1 | ENSG00000114115 | 8 |
| PNLIPRP3 | ENSG00000203837 | 8 |
| KLRC1 | ENSG00000134545 | 8 |
| KLRD1 | ENSG00000134539 | 8 |
| ZBTB16 | ENSG00000109906 | 8 |
| NKG7 | ENSG00000105374 | 5 |
| IL1RL1 | ENSG00000115602 | 8 |
| LIF | ENSG00000128342 | 8 |
| LRP4 | ENSG00000134569 | 8 |
| PTGS1 | ENSG00000095303 | 8 |
| TPSB2 | ENSG00000197253 | 6 |
| MLANA | ENSG00000120215 | 8 |
| PMEL | ENSG00000185664 | 8 |
| TYR | ENSG00000077498 | 8 |
| KIT | ENSG00000157404 | 8 |
| ALDH2 | ENSG00000111275 | 8 |
| CD9 | ENSG00000010278 | 8 |
| MSR1 | ENSG00000038945 | 8 |
| S100A11 | ENSG00000163191 | 6 |
| S100A9 | ENSG00000163220 | 5 |
| TREM1 | ENSG00000124731 | 8 |
| VSIG4 | ENSG00000155659 | 8 |
| S100A8 | ENSG00000143546 | 4 |
| COL24A1 | ENSG00000171502 | 8 |
| CRABP1 | ENSG00000166426 | 5 |
| ITGA11 | ENSG00000137809 | 8 |
| MMP23B | ENSG00000189409 | 4 |
| PTGFR | ENSG00000122420 | 8 |
| TNN | ENSG00000120332 | 8 |
| CEBPA | ENSG00000245848 | 8 |
| CLCA4 | ENSG00000016602 | 8 |
| CYP1A1 | ENSG00000140465 | 8 |
| LYPD3 | ENSG00000124466 | 8 |
| MYCL | ENSG00000116990 | 8 |
| LCE5A | ENSG00000186207 | 7 |

| | | |
|---|---|---|
| ACOX2 | ENSG00000168306 | 8 |
| AWAT2 | ENSG00000147160 | 8 |
| CYP4F8 | ENSG00000186526 | 8 |
| DGAT2 | ENSG00000062282 | 8 |
| FADS1 | ENSG00000149485 | 8 |
| KRT79 | ENSG00000185640 | 8 |
| DBX2 | ENSG00000185610 | 8 |
| PLA2G3 | ENSG00000100078 | 8 |
| RDH12 | ENSG00000139988 | 8 |
| SUSD2 | ENSG00000099994 | 8 |
| CDH3 | ENSG00000062038 | 8 |
| CDT1 | ENSG00000167513 | 8 |
| DLK2 | ENSG00000171462 | 8 |
| GDPD2 | ENSG00000130055 | 8 |
| KREMEN2 | ENSG00000131650 | 8 |
| KRT31 | ENSG00000094796 | 8 |
| LAMB4 | ENSG00000091128 | 8 |
| SYT8 | ENSG00000149043 | 7 |
| CFTR | ENSG00000001626 | 8 |
| FOXA1 | ENSG00000129514 | 8 |
| GRIA1 | ENSG00000155511 | 8 |
| NOTUM | ENSG00000185269 | 8 |
| RHCG | ENSG00000140519 | 8 |
| SCGB2A2 | ENSG00000110484 | 8 |
| TMPRSS2 | ENSG00000184012 | 8 |
| GPR20 | ENSG00000204882 | 8 |
| KCNA5 | ENSG00000130037 | 8 |
| PDE4C | ENSG00000105650 | 8 |
| RERGL | ENSG00000111404 | 8 |
| TBX2 | ENSG00000121068 | 8 |
| COL5A3 | ENSG00000080573 | 8 |
| GJC1 | ENSG00000182963 | 8 |
| MUC3A | ENSG00000169894 | 8 |
| PDE5A | ENSG00000138735 | 8 |
| STEAP4 | ENSG00000127954 | 8 |
| TRPC6 | ENSG00000137672 | 8 |
| ACTA1 | ENSG00000143632 | 8 |
| ACTG2 | ENSG00000163017 | 8 |
| GPR183 | ENSG00000169508 | 8 |

| | | |
|---|---|---|
| MYOCD | ENSG00000141052 | 8 |
| CSPG4 | ENSG00000173546 | 8 |
| DES | ENSG00000175084 | 8 |
| RGS5 | ENSG00000143248 | 8 |
| CD68 | ENSG00000129226 | 8 |
| FCGR1A | ENSG00000150337 | 8 |
| FCGR3A | ENSG00000203747 | 8 |
| CD14 | ENSG00000170458 | 8 |
| GALR1 | ENSG00000166573 | 8 |
| NRXN1 | ENSG00000179915 | 8 |
| SOX10 | ENSG00000100146 | 8 |
| SOX2 | ENSG00000181449 | 8 |
| ZNF536 | ENSG00000198597 | 8 |
| CCL1 | ENSG00000108702 | 8 |
| CCL2 | ENSG00000108691 | 8 |
| CCL11 | ENSG00000172156 | 8 |
| CCL13 | ENSG00000181374 | 8 |
| CCL24 | ENSG00000106178 | 6 |
| CXCL1 | ENSG00000163739 | 8 |
| CXCL16 | ENSG00000161921 | 8 |
| CXCL2 | ENSG00000081041 | 8 |
| CXCL9 | ENSG00000138755 | 8 |
| CCL4 | ENSG00000275302 | 8 |
| CCL5 | ENSG00000271503 | 8 |
| CXCL5 | ENSG00000163735 | 8 |
| CXCL13 | ENSG00000156234 | 8 |
| CCL18 | ENSG00000275385 | 8 |
| C4B | ENSG00000224389 | 8 |
| C5 | ENSG00000106804 | 8 |
| C5AR1 | ENSG00000197405 | 8 |
| CFB | ENSG00000243649 | 8 |
| F2 | ENSG00000180210 | 8 |
| F3 | ENSG00000117525 | 8 |
| IL37 | ENSG00000125571 | 8 |
| IFNAR1 | ENSG00000142166 | 8 |
| IFNGR1 | ENSG00000027697 | 8 |
| IL13RA1 | ENSG00000131724 | 8 |
| IL13RA2 | ENSG00000123496 | 8 |
| IL17RA | ENSG00000177663 | 8 |

| | | |
|---|---|---|
| IL17RB | ENSG00000056736 | 8 |
| IL18R1 | ENSG00000115604 | 8 |
| IL1R1 | ENSG00000115594 | 8 |
| IL1R2 | ENSG00000115590 | 8 |
| IL2RB | ENSG00000100385 | 8 |
| IL31RA | ENSG00000164509 | 8 |
| IL12A | ENSG00000168811 | 8 |
| IL18 | ENSG00000150782 | 8 |
| IL1B | ENSG00000125538 | 8 |
| IL31 | ENSG00000204671 | 8 |
| IL32 | ENSG00000008517 | 8 |
| IL36G | ENSG00000136688 | 8 |
| IL5 | ENSG00000113525 | 8 |
| IL6 | ENSG00000136244 | 8 |
| IL7 | ENSG00000104432 | 8 |
| CSF2 | ENSG00000164400 | 7 |
| CSF3 | ENSG00000108342 | 8 |
| EREG | ENSG00000124882 | 8 |
| TSLP | ENSG00000145777 | 8 |
| VEGFA | ENSG00000112715 | 8 |
| WNT4 | ENSG00000162552 | 8 |
| WNT5B | ENSG00000111186 | 8 |
| WNT10A | ENSG00000135925 | 8 |
| WNT11 | ENSG00000085741 | 8 |
| DKK1 | ENSG00000107984 | 8 |
| DKK3 | ENSG00000050165 | 8 |
| CHRD | ENSG00000090539 | 8 |
| BMP1 | ENSG00000168487 | 8 |
| BMP2 | ENSG00000125845 | 8 |
| BMP4 | ENSG00000125378 | 8 |
| BMP6 | ENSG00000153162 | 8 |
| BMP7 | ENSG00000101144 | 8 |
| CTNNB1 | ENSG00000168036 | 8 |
| DLL1 | ENSG00000198719 | 8 |
| DLL4 | ENSG00000128917 | 8 |
| GLI1 | ENSG00000111087 | 8 |
| NOTCH1 | ENSG00000148400 | 8 |
| SHH | ENSG00000164690 | 8 |
| GLI2 | ENSG00000074047 | 8 |

| | | |
|---|---|---|
| ACVRL1 | ENSG00000139567 | 8 |
| IGF1 | ENSG00000017427 | 8 |
| IGFBP4 | ENSG00000141753 | 8 |
| INHBA | ENSG00000122641 | 8 |
| INHBB | ENSG00000163083 | 8 |
| VEGFC | ENSG00000150630 | 8 |
| LTBP1 | ENSG00000049323 | 8 |
| EBI3 | ENSG00000105246 | 8 |
| GLT8D2 | ENSG00000120820 | 8 |
| HPSE2 | ENSG00000172987 | 8 |
| FAM83A | ENSG00000147689 | 8 |
| VASH1 | ENSG00000071246 | 8 |
| CD1E | ENSG00000158488 | 8 |
| EMID1 | ENSG00000186998 | 8 |
| PCP4 | ENSG00000183036 | 7 |
